# RBM20-variants induce distinct calcium handling and metabolic phenotypes in patient-specific stem cell models of dilated and non-compaction cardiomyopathy

**DOI:** 10.1101/2025.01.13.632728

**Authors:** Sabine Rebs, Farbod Sedaghat-Hamedani, Elham Kayvanpour, Jan Dudek, Branimir Berecic, Hanna Eberl, Daniela Hubscher, Christoph Reich, Teresa Klein, Soren Doose, Viacheslav O Nikolaev, Kaomei Guan, Gerd Hasenfuss, Markus Sauer, Malte Tiburcy, Christoph Maack, Benjamin Meder, Katrin Streckfuss-Bomeke

## Abstract

**Background and aim:** Mutations in the splice regulator RBM20 account for ∼3 % of genetic cardiomyopathies. In particular, the highly conserved RS domain is a hotspot for disease-associated mutations. Previously, mutations at same amino acid position 634 in the hotspot RS-domain were found to cause dilated cardiomyopathy (DCM) with left ventricular non-compaction (R634L) or without (R634W), but the pathophysiological mechanisms that govern the heterogeneity in phenotype presentation remained unknown. Here, we identify the molecular events caused by the distinct RBM20 mutations from DCM and left-ventricular non-compaction (LVNC) using patient-specific stem cell models.

**Methods:** We generated induced pluripotent stem cell-derived cardiomyocytes (iPSC-CM) of one LVNC- and two DCM-patients harboring the RBM20-mutations R634L (LVNC) or R634W (DCM). We investigated alternative splicing activity, RBM20 localization, sarcomeric regularity, cAMP level, kinase-specific phosphorylation of key Ca^2+^ handling enzymes, physiological cardiac functions as Ca^2+^ homeostasis, and metabolic activity on a patient-specific cardiomyocyte level. Force generation was analyzed in patient-specific engineered myocardial tissues. Isogenic rescue and mutation insertion lines were generated by CRISPR/Cas9 technology to analyze the direct impact of the RBM20 mutations on the cardiac phenotype.

**Results:** We observed common splicing aberrations for LVNC- and DCM-CM in *TTN* and *RYR2*, RBM20 cytoplasmatic accumulation and irregular sarcomeric structure. LVNC-CM harboring the RBM20-p.R634L variant show distinct molecular, cellular and functional impairments that manifest in *CAMK2D*, *TRDN* and *IMMT* mis-splicing. Splicing defects in LVNC-CM correlate with elevated systolic Ca^2+^ and faster Ca^2+^ kinetics with elevated cAMP levels and PLN-hyperphosphorylation. An increased metabolic activity and mitochondrial membrane potential support the ‘hyperactive’ LVNC-CM. By contrast, DCM-CM (RBM20-p.R634W) distinctly present with decreased systolic Ca^2+^ and increased SR Ca^2+^leak but unchanged Ca^2+^ kinetics and metabolic activity. Both mutations lead to severely reduced force of contraction in engineered myocardium. CRISPR/Cas9 gene-edited isogenic control lines of both described RBM20 mutations in LVNC and DCM demonstrated the causative nature of the two mutations and their diverging effects. Further, L-type Ca^2+^ channel blockade by verapamil ameliorates the Ca^2+^ cycling and leakage phenotypes in LVNC- and DCM-CM.

**Conclusion:** We show the first comparative iPSC-model of splice-defect-associated RBM20-dependent LVNC-p.R634L and DCM-p.R634W. We found shared and variant-specific phenotypes on a patient-specific level. Our data demonstrate that the different RBM20 mutations manifest in distinct molecular aberrations in alternative splicing and RBM20 cytoplasmic accumulation that convey various physiological impairments in structure, Ca^2+^ handling, metabolism and contractile force.

## Introduction

Heart failure is among the most common causes of hospitalization in Europe and the United States and is associated with high morbidity and mortality. Among patients with heart failure with reduced ejection fraction (HFrEF), non-ischemic dilated cardiomyopathy (DCM) contributes to 30-40 % of cases. DCM is characterized by left ventricular (LV) dilation, thinning and contractile dysfunction [1, 2]. In contrast, left-ventricular non-compaction cardiomyopathy (LVNC) is a rarer cardiomyopathy, which is characterized by a hyper-trabeculated myocardium showing deep recesses within the ventricular muscle and can additionally be accompanied by dilation of the chambers [2]. LVNC is either the result of a developmental defect with incomplete myocardial compaction that generally occurs during the 5^th^ to 8^th^ week of embryogenesis or later in life due to genetic defects or afterload increases [3]. Albeit having distinct clinical features, the recent European guidelines classify non-compaction or hyper-trabeculation as an additional phenotypic trait of DCM, rather than a distinct cardiomyopathy (ESC 2023 Management of Cardiomyopathies guideline). Numerous genetic studies have revealed overlapping gene loci for LVNC and DCM, including many that affect sarcomeric or structural proteins, some having a severely impaired prognosis [4–6]. Hence, genetic testing has a pivotal role in diagnosis and risk stratification in both patient groups [7]. Mutations in the RNA binding motif protein 20 (RBM20) underlie 2-6 % of all genetic DCM cases [8]. In general, RBM20-based DCM presents as highly penetrant and arrhythmogenic with a high risk for sudden cardiac death [9]. In 2017, a clinical study associated the distinct RBM20-variant R634L with LVNC, which has not been observed in DCM cases [6]. RBM20 is a major cardiac splicing factor with more than 30 target mRNAs linked to cardiac muscle function such as *RYR2*, *TTN,* or *CAMK2D*. The mutational hotspot within the RBM20 gene is located at the highly conserved RS-domain (p.633-638 - PRSRSP) in exon 9, which is crucial for protein-protein interaction with the spliceosome and for nuclear localization upon phosphorylation of serines. In this hotspot, 16 mutations have been described as pathogenic variants, including the DCM-variant p.R634W and the LVNC-variant p.R634L [10]. Recent studies expanded the pathogenic scope of RBM20 mutations to hypertrophic cardiomyopathy [11, 12]. There is also evidence that mutations outside the RS-domain are also causative for inherited DCM [13]. However, if and how mutations in the same gene and even the same amino acid position can determine different cardiomyopathies needs a deeper mechanistic understanding. The generation of induced pluripotent stem cells (iPSC) derived from patients, as well as the introduction of mutations by CRISPR/Cas9 genome editing, is an invaluable tool in the research field of precision medicine [14, 15]. The successful differentiation of functional iPSC-cardiomyocytes (CM) was an important step in cardiac research and many studies have hence demonstrated that patient-specific iPSC-CM recapitulate disease-related cardiac phenotypes *in vitro* [16–18]. Although RBM20 research has spanned the model organisms of mice, rats, and pigs, the animal models often represent an RBM20 knockout model or a homozygous variant rather than the clinically relevant heterozygous missense mutations or have a species-specific limitation of animal physiology [19–21]. Of note, very rare heterozygous loss of function variants were reported recently in the context of arrhythmia [22]. Some studies used patient-specific-iPSC for analysis of RBM20 mutations (p.S635A and R636S) [17, 23] while others have introduced a patient-relevant mutation into a healthy iPSC line (P633L, R634Q, R636S) [24, 25]. In general, the pathological effects of RBM20 mutations are attributed to erroneous splicing of target genes and/or RBM20 protein accumulation in the cytoplasm. However, it is under debate, which one of these two is the predominant driver for the development of cardiomyopathies. Variants in RBM20 generally harm the sarcomeric structure and cardiac Ca^2+^ cycling, resulting in reduced contractile performance [10].

Here, we analyzed two genetic variants in RBM20 at the very same amino acid position p.R634, which were linked to either LVNC (p.R634L) or DCM phenotype (p.R634W), and investigated the pathogenesis of these mutations on molecular, cellular, and functional levels. We used iPSC-CM generated from respective cardiac patients and compared them to isogenic rescue and insertion lines generated by CRISPR/cas9 genome editing. A comparable analysis of both RBM20 variants using patient-specific stem cell models allows for an in-depth analysis of RBM20-linked LVNC and DCM. Furthermore, this study expands the scope of RBM20-pathology into metabolic impairments and preclinical therapeutic interventions with Ca^2+^ channel inhibitors.

## Material & Methods

Details are available in the supplements.

## Results

### Patient-specific iPSC-CM from two different cardiomyopathy families with distinctive RBM20 mutations

To study the relationship between RBM20 variants in the context of two different cardiomyopathy traits, we recruited one patient diagnosed with LVNC (DCM with non-compaction trait) and two patients with DCM (without non-compaction), each from different families (Fig. 1A), for the reprogramming of somatic material into iPSC lines. The LVNC index patient (LVNC pedigree II.3) harbors the RBM20 variant p.R634L and was diagnosed with a severe form of LVNC and heart failure [6]. Diagnosed at age 39, she had a LVEF of only 20 %, which led to the implantation of an implantable cardioverter-defibrillator (ICD). Her three children are also affected by heart disease: III.1 died at a young age due to tetralogy of Fallot. The other son and daughter both inherited the RBM20 variant p.R634L with varying severities. Her daughter (III.3) exhibits LV hyper-trabeculation but is symptom-free with a normal LVEF (56 %) at age 35. The son (III.2) underwent heart transplantation at age 17 due to severe LVNC and heart failure. The RBM20 DCM-linked variant p.R634W was initially identified in cohort screenings [8]. Two brothers (DCM pedigree III.8, III.10) characterized with DCM and reduced LVEF (40 % and 30 % respectively) harbor this RBM20 variant p.R634W. Additionally, the daughter (IV.4) of DCM1 (III.8) inherited the p.R634W variant and also suffers from DCM (Fig. 1A). The two DCM patients are presented separately by different colors or symbols. To evaluate the pathogenic potential of p.R634L and p.R634W mutations, patient-specific RBM20 iPSC were generated. CRISPR/Cas9 was applied to produce isogenic rescue iPSC lines, termed resLVNC and resDCM1 (Fig. 1A). In addition to healthy controls previously described by us [16], two healthy family members from the DCM family (II.3 and III.3) donated somatic material (Fig. 1A). The generation and characterization of DCM-iPSC lines from patient III.8 (here termed DCM1) and the corresponding isogenic control line (here termed resDCM1) was recently published by our groups [26]. The remaining iPSC lines generated in this study exhibit full pluripotency and spontaneous *in vitro* differentiation capacity (Suppl. Fig. 1). Furthermore, the patient lines show genetic integrity even after gene editing (Suppl. Fig. 2A) and harbor the expected nucleotide sequences (Suppl. Fig. 2B). All iPSC lines successfully differentiated into beating cardiomyocytes (iPSC-CM) with an efficiency of 80 – 97 % cardiac troponin T (cTNT)-positive cells (Suppl. Fig. 2C) and high expression of the ventricular marker *MYL2* concomitant with low expression of the atrial marker *NR2F2* (Suppl. Fig. 2D). Taken together, the generated iPSC-lines provide a suitable model for the study of RBM20-linked cardiomyopathies.

**Fig. 1:**
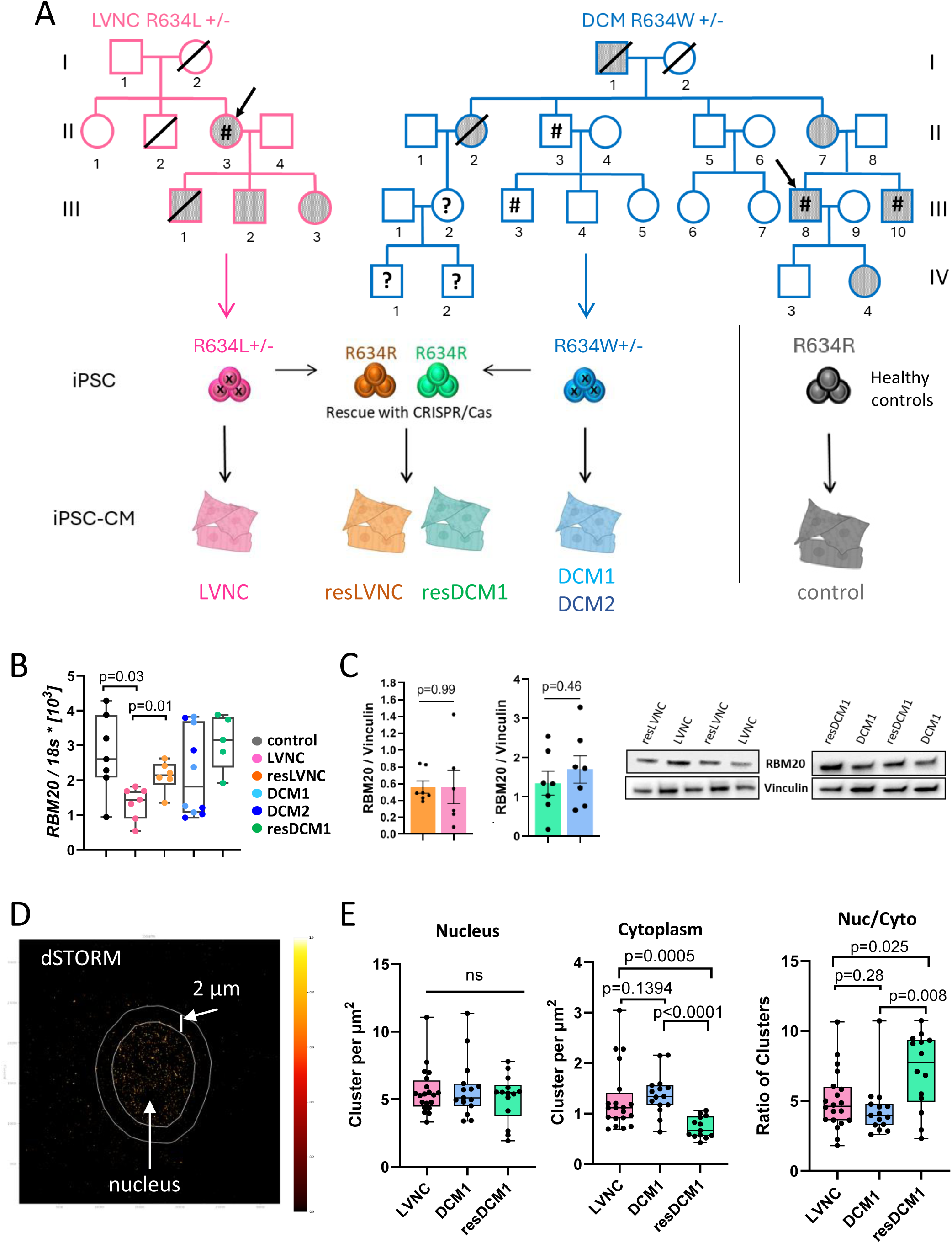
Generation of iPSC-CM of LVNC- and DCM-affected family members with RBM20 mutations. **A:** Pedigree of LVNC- and DCM-affected families carrying a RBM20 mutation at amino acid position R634. Filled symbols labeled disease-affected individuals. Symbols marked with a (#) are individuals who donated somatic material used for reprogramming into iPSC. Crossed symbols mean the individual is deceased. The arrow marks the patient of which isogenic control lines were derived. The control category (grey/black) comprises iPSC-CM, which was derived from healthy DCM family members and unrelated healthy individuals already published by our group. **B:** QPCR analysis of *RBM20*-mRNA expression in control and patient iPSC-CM. Each dot represents one differentiation experiment. Data is presented as box plots. P-values were determined by Kruskal-Wallis test against control with Dunn,s correction. The Mann-Whitney test was conducted between disease and respective rescue-line (p-values are marked with #). **C:** Protein quantification of RBM20. Example of Western blot membranes for RBM20 and Vinculin (right) and quantification of RBM20 resLVNC vs LVNC and resDCM1 vs DCM (left). P-values by Mann-Whitney test. **D:** Representative RBM20 *d*STORM image from DCM-CM. Two ROIs per cell, nucleus, and cytoplasm, were analyzed with LOCAN. **E:** Quantification of RBM20 cluster densities, calculated for the nucleus (Nuc), the cytoplasm (Cyto), and the ratio between Nuc to Cyto obtained by *d*STORM imaging and LOCAN analysis. Each dot in the box plot represents one cell. P-values calculated by Mann-Whitney test.

### RBM20 mutations affect protein localization and RBM20-dependent splice target

The analysis of *RBM20* mRNA levels in the generated patient-specific iPSC-CM revealed that LVNC-CM, but not DCM-CM showed significantly decreased *RBM20* mRNA levels, which is rescued in the isogenic resLVNC-CM (Fig. 1B). However, this effect was not observed for RBM20 protein levels (Fig. 1C). Since the mutations are located in the RS-domain, which is essential for nuclear localization, we analyzed the impact of both variants on the localization of the RBM20 protein. Immunofluorescence stainings of RBM20 demonstrated that for both, LVNC- and DCM-CM, RBM20 signals are detected in the sarcoplasm and the nucleus, whereas for healthy and isogenic iPSC-CM the RBM20 signal is distinctly limited to the nucleus (Suppl. Fig. 3A). To quantify this observation super-resolution imaging of RBM20 was performed using single-molecule localization microscopy by *direct* stochastic optical reconstruction microscopy (*d*STORM) [27, 28] (Fig. 1D). Due to technical limitations only one isogenic rescue line served as a control. RBM20 was stained with Alexa Fluor 647 by indirect immunofluorescence. Employing customized software LOCAN [29] with a DBSCAN [30] clustering algorithm we observed that the ratio of RBM20 densities (nucleus to sarcoplasm) in LVNC- and DCM-CM is significantly reduced compared to the isogenic rescue line resDCM1 (Fig. 1D, E). The results indicate that RBM20 protein aberrantly accumulates in the sarcoplasm if there is a missense mutation at position p.634, regardless of the amino acid switch to either Leu or Trp (LVNC or DCM respectively). Taken together, while the two RBM20 mutations do not affect RBM20 protein expression, both variants cause RBM20 to aberrantly accumulate in the sarcoplasm instead of the nucleus.

Besides localization, we assessed the effects of RBM20 mutations p.R634L and p.R634W on RBM20-dependent splice targets. RBM20 splice targets have been well defined since the first description by Guo and colleagues in 2012 [20] and can be categorized functionally into Ca^2+^ handling genes, ion channels and structural proteins/genes. Here, we selected splice targets from RBM20 knockout rats and translated them to the human loci [31]. Splicing of titin (*TTN*) and ryanodine receptor 2 (*RYR2*) demonstrated RBM20-dependent mis-splicing in LVNC- and DCM-CM (Figure 2A). The *TTN* mRNA showed transcripts with an increased ratio of longer (N2BA) to shorter (N2B) isoform due to decreased N2B levels (Suppl. Fig. 3B). The shorter N2B isoform is increased during maturation and annotated as the adult *TTN* form. Expression of *RYR2*-mRNA that includes a cryptic 24bp insertion from intron 80-81 was also increased for LVNC- and DCM-CM compared to control and rescue lines (Fig. 2A). By contrast, transcripts with included exon 5 in LIM domain binding 3 (*LDB3)* was only increased in DCM-CM (Fig. 2B). Furthermore, the inclusion of exon 9 in triadin (*TRDN)* as well as inclusion of a nuclear localization signal (NLS) in exon 14 of the Ca^2+^/calmodulin-dependent protein kinase type 2 delta (*CAMK2D)* and exon5/6 of inner mitochondrial membrane protein (*IMMT)* were only affected in LVNC-CM, but not in DCM-CM compared to control-CM (Fig. 2C). These splicing defects were rescued in the resLVNC- and resDCM1-CM demonstrating that the individual mutations are the driver of molecular alterations (Fig. 2A-C). In conclusion, these data suggest that splicing of RBM20-targets was affected differentially depending on the missense mutation R634L or W, while mis-localization of the RBM20-protein was similar for both missense variants. To gain further mechanistic insight, gene expression and splice isoform profiling were performed. Principal component analysis (PCA) plot shows clustering of disease versus rescue lines, but some overlap between DCM and LVNC (Suppl. Fig. 3C). We identified 1874 genes that are differentially expressed between resLVNC and LVNC, with 718 downregulated and 1156 upregulated (adjusted p-value <= 0.05, abs (fold change) > 1.5) and 3377 genes between resDCM1 and DCM with 1782 downregulated and 1595 upregulated (adjusted p-value <= 0.05, abs (fold change) > 1.5). Only 24 genes were differentially expressed (adjusted p-value <= 0.05, abs (fold change) > 1.5) between LVNC and DCM, whereas 14 of these genes account for y-linked genes since DCM lines are derived from a male patient (Suppl. Fig. 3D). Notably, gene expression of RBM20 splice targets is scarcely affected in LVNC- and DCM-CM (Suppl. Fig. 3E). Furthermore, gene ontology (GO) analysis of the top 100 differentially expressed genes for resLVNC vs LVNC and for resDCM1 vs DCM1 does show only a few cardiac relevant GO term hits (Suppl. Fig. 4A/B, Suppl. Tab. 7/8). Taken together, this suggests that gene expression differences are unlikely to drive the disease phenotypes observed for LVNC- and DCM-CM. However, analyzing the gene expression data for gene splice isoform expression, we observed that known RBM20 splice targets are spliced differently depending on the RBM20-variant. Intriguingly, only two of the 36 RBM20 splice targets analyzed are identically spliced between LVNC- and DCM-CM, which are *MYH7* and *TTN* (Fig. 2D), while others are distinct for LVNC such as *SH3KPB1* or *NEXN* for DCM-CM (Suppl. Fig. 5).

**Fig. 2:**
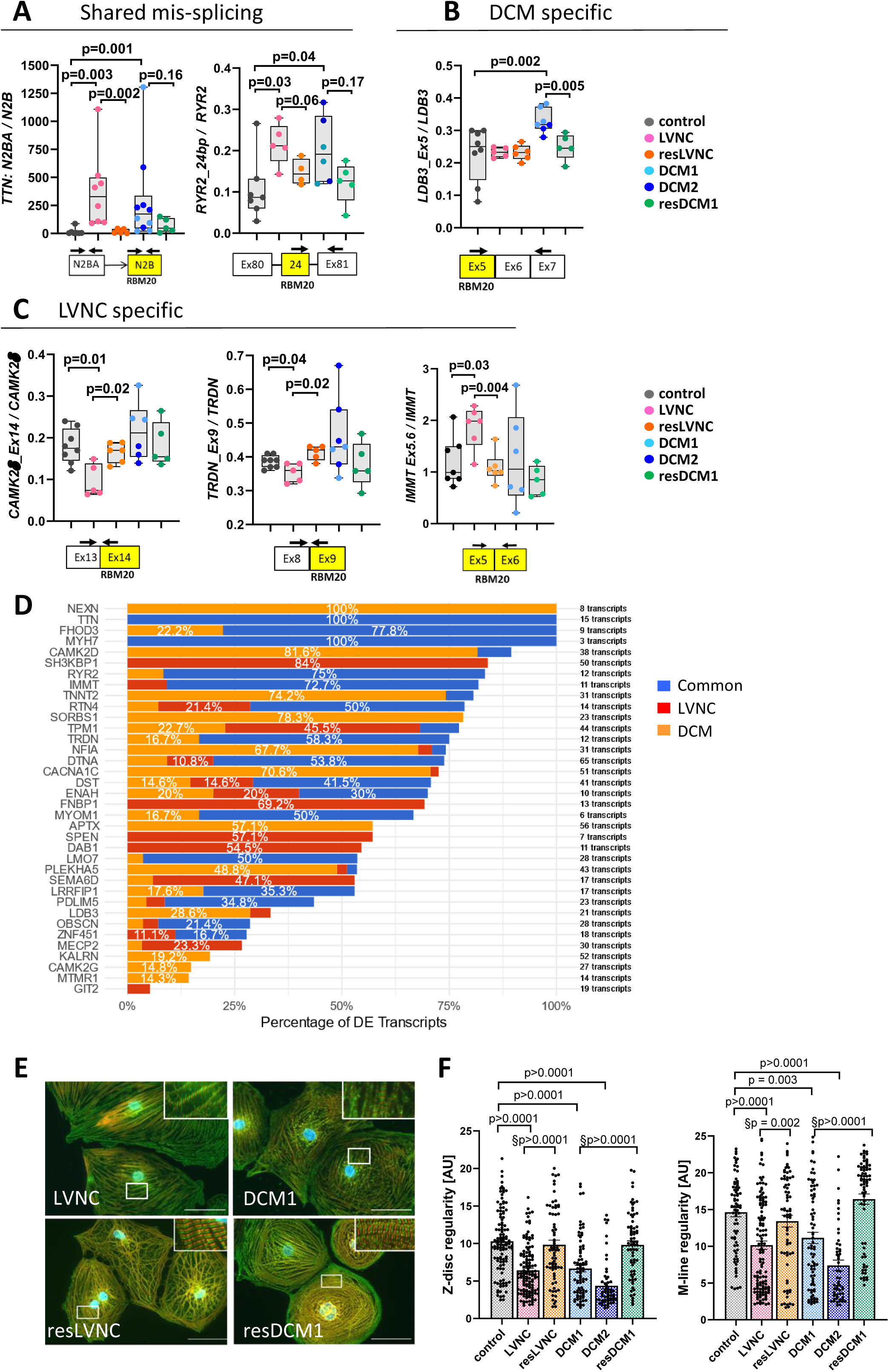
RBM20-dependent splice and sarcomeric defects in LVNC- and DCM-CM. **A-C:** QPCR analysis of RBM20 splice targets. Data is shown as box plots, whereas every dot represents one differentiation experiment. P-values were determined by Mann-Whitney test control vs. patient line (LVNC, DCM1, DCM2) and patient line vs. rescue line (resLVNC, resDCM1). **A:** Shared mis-splicing events in LVNC- and DCM-CM. The primers were designed against the N2BA and N2B domain of *TTN*, and an intronic 24 bp insertion of *RYR2*. **B-C:** Differential mis-splicing for exon 14 in *CAMK2D* and exon 9 in *TRDN* for LVNC-CM and exon 5 in *LDB3* in DCM-CM. **D:** The stacked bar chart depicts the differentially expressed exon usage percentage across a selection of RBM20 target genes. For each gene, the total number of unique transcripts is shown as a percentage of three categories: common differentially expressed (DE) exon usage for DCM and LVNC (blue), DE exon usage unique to DCM (orange), and DE exon usage unique to LVNC (red). The y-axis lists RBM20 target genes. The numbers on the bars indicate the actual percentages for segments greater than 10 %. In addition, the total number of unique transcripts per gene is shown to the right of the bars. **E:** Representative immunofluorescence stainings of LVNC-, resLVNC-, DCM-, and resDCM1-CM. Visualization of sarcomeric Z-disc by staining against α-actinin (green) and the M-line by TTN (M8/M9 antibody) (red). Scale bars: 50 µm. **F:** Quantifying Z-disc (α-actinin) and M-line (TTN) sarcomeric regularity using Fast Fourier transformation. Bar graphs depict the peak amplitude of the first-order peak. Multiple pictures were analyzed [number of differentiations/analyzed pictures] for control [6/112], LVNC [7/128], resLVNC [4/73], DCM1 [4/79], DCM2 [3/58] and resDCM1 [4/69]. Data is represented as a bar graph with mean +/-SEM. P-values were determined by Kruskal-Wallis test against control with Dunn,s correction. Additionally, the patient- and corresponding isogenic rescue lines were analyzed using Mann-Whitney test (p-values marked with §).

In conclusion, the gene profiling revealed that differential gene isoform expression of RBM20 target genes is a more dominant disease driver for these RBM20-dependent cardiomyopathies instead of the change in the overall gene expression.

### Sarcomeric regularity is impaired in LVNC- and DCM-CM

A sarcomere is the smallest contractile unit of a CM, and the amount and regularity of the sarcomeres are responsible for contractile properties of cardiac cells [32]. Since both RBM20 mutations R634L (LVNC) and R634W (DCM) alter the *TTN* isoform, we focused on the sarcomeric organization pattern in iPSC-CM. To this end, double staining was used to visualize the sarcomeric Z-disc with α-actinin and the M-line with TTN M8/M9 antibodies (Fig. 2E) followed by quantification of sarcomeric regularity using fast Fourier Transformation (FFT). We observed a significant impairment in sarcomeric regularity for the Z-disc and M-line in LVNC-CM as well as DCM-CM compared to control-CM (Fig. 2F). Isogenic rescue iPSC-CM showed a regular sarcomeric structure comparable to the control cells, demonstrating the importance of wt-RBM20 for sarcomeric regularity. As sarcomeric irregularity is a common trait for RBM20 mutations, we cannot distinctly pinpoint whether this is due to *TTN* mis-splicing or RBM20 mis-localization as these molecular aberrations are shared between LVNC- and DCM-CM.

### Dysfunctional Ca^2+^ homeostasis and contractility for LVNC- and DCM-CM

Cardiac contractility is governed by the processes of excitation-contraction (EC) coupling, where Ca^2+^ handling plays a key role [33]. Since we have shown that RBM20 affects the splicing of Ca^2+^ handling genes such as *RYR2, TRDN* or *CAMK2D*, we evaluated whether the different RBM20 mutations affect Ca^2+^ homeostasis. Employing the cytosolic Ca^2+^-dye Fluo-4-AM, we observed accelerated rates of Ca^2+^ rise and decay for LVNC and DCM (i.e., shorter times to peak or 50 % decay, respectively) with LVNC-CM showing the highest absolute decrease (Fig. 3A/B, Suppl. Fig. 6A). The most important regulator of cardiac contractility is β-adrenergic stimulation and therefore, we treated the CM with β-adrenergic agonist Isoprenaline (Iso). While control- and DCM-CM demonstrated a prominent acceleration in rise time (time to peak) after Iso treatment, this was not the case in LVNC-CM, where the basal rates were already pre-accelerated (Fig. 3B). This pre-acceleration was rescued in resLVNC-CM, which thereby also restored the β-adrenergic response.

**Fig. 3:**
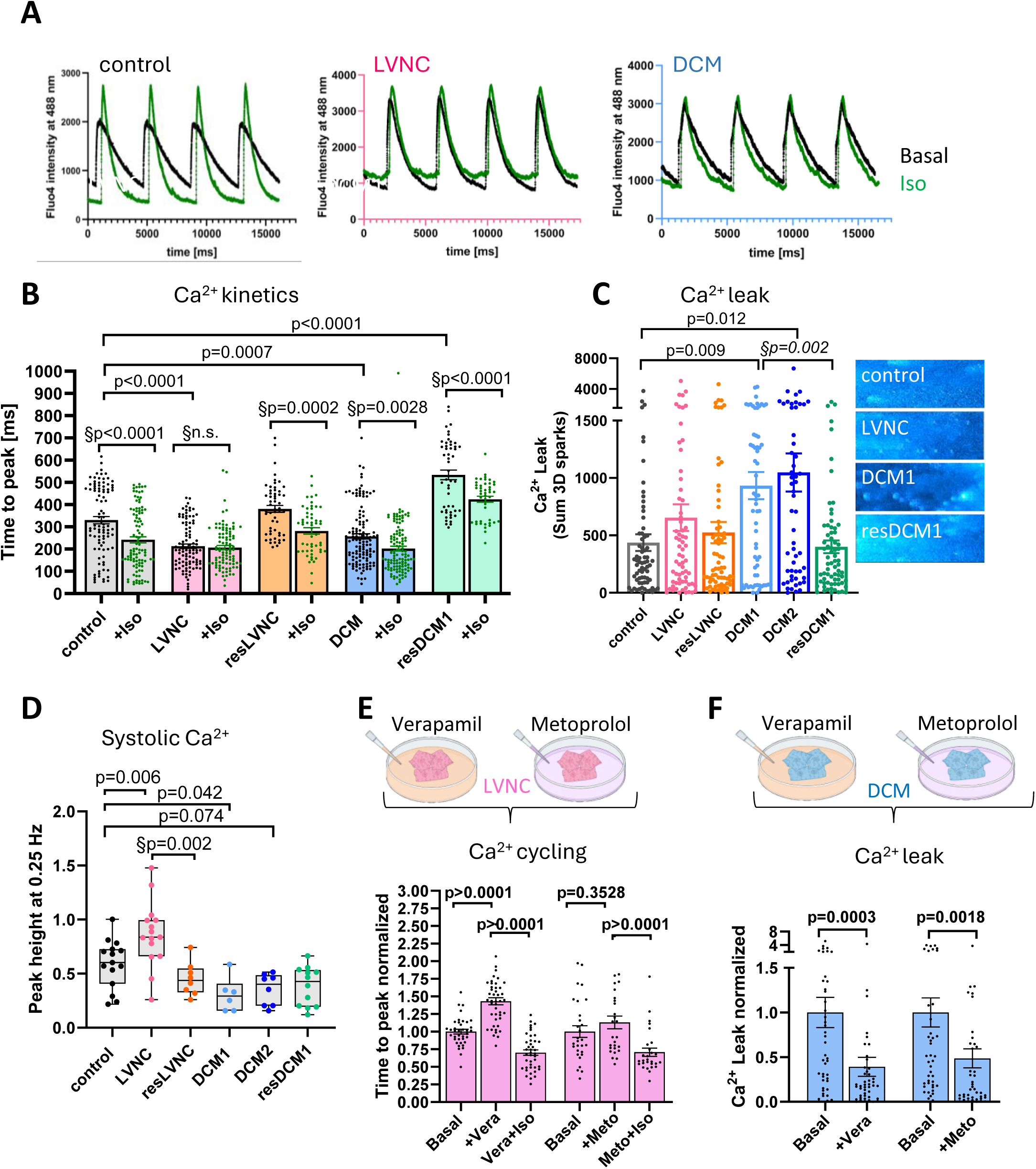
LVNC- and DCM-CM show differential Ca^2+^ handling pathologies. **A:** Exemplary single-cell Ca^2+^ traces from iPSC-CM of control, LVNC and DCM at basal (black) and Isoprenaline (Iso) stimulation (1 µM) measured with Fluo-4-AM. **B:** Ca^2+^ transient rise time (time to peak) at basal and Iso-stimulated (1 µM) conditions. LVNC-CM showed decreased basal rise time levels and no measurable reaction to Iso (non significant = n.s.). [number of differentiations/analyzed cells] for basal: control [6/94], LVNC [6/101], resLVNC [3/55], DCM [7/126] and resDCM1 [3/51] and Iso stimulated: control [6/101], LVNC [6/97], resLVNC [3/53], DCM [7/119] and resDCM1 [3/44]. Data is represented as bar graph with mean +/-SEM. P-value by Kruskal-Wallis test against control (basal values) with Dunn,s correction. §P-values by Two-way ANOVA Sidak,s multiple comparison test Basal vs Iso. **C:** Increased Ca^2+^ leakage in DCM-CM. Quantification of Ca^2+^ leak (left) and representative original confocal line scans showing diastolic SR Ca^2+^ sparks in iPSC-CM using the ImageJ sparkmaster plugin (right). Quantitative calculation of 3D diastolic SR Ca^2+^ leakage for: [number of differentiations/analyzed cells] control [4/70], LVNC [4/67], resLVNC [4/70], DCM1 [4/63], DCM2 [3/55] and resDCM1 [4/71]. Data is represented as bar graph with mean +/-SEM. P-values were calculated by Kruskal-Wallis test against control with Dunn,s correction. §P-value was calculated with Mann-Whitney test: patient vs respective isogenic line. **D:** LVNC-CM showed elevated, and DCM-CM reduced systolic Ca^2+^. Measurements with ratiometric dye Fura-2-AM. [number of differentiations/analyzed cells] for control [5/15], LVNC [5/15], resLVNC [2/8], DCM1 [2/6] and DCM2 [3/12]. P-values were calculated by One-way ANOVA with Dunnett,s correction against control. §P-value was calculated with Mann-Whitney test: patient vs respective isogenic line. **E:** Ca^2+^ kinetics in LVNC-CM after drug treatments. P-value was calculated by Mann-Whitney test. [number of differentiations/analyzed cells] for Verapamil set: Basal – Vera – Vera+Iso [3/38 – 3/45 – 3/41]. Metoprolol set: Basal – Meto – Meto+Iso [2/29 – 2/27 – 2/28]. **F:** Ca^2+^ sparks analysis in DCM-CM after drug treatment. P-value was calculated by Mann-Whitney test. [number of differentiations/analyzed cells] for Basal – Vera [3/44 – 3/45] and Basal – Meto [3/47 – 3/45]. Vera: verapamil; Meto: metoprolol

Diastolic SR Ca^2+^ release via leaky RYR2 is a key mechanism for contractile dysfunction and arrhythmias in HF and other cardiac diseases [34]. We observed that the Ca^2+^ leakage was significantly increased in DCM-CM of both patients (DCM1 and 2), but not LVNC-CM, which was reversed in resDCM1-CM (Fig. 3C). Furthermore, we investigated the impact of RBM20-mutations on systolic Ca^2+^ transients, SR Ca^2+^ load, and SR fractional release using the ratiometric Ca^2+^-dye Fura-2-AM. LVNC-CM had increased systolic Ca^2+^ transient amplitudes, whereas DCM-CM exhibited reduced transients compared to control-CM (Fig. 3D). There were no significant changes in SR Ca^2+^ load or fractional SR Ca^2+^ release, although DCM-CM tended to have reduced SR Ca^2+^ load (Suppl. Fig. 6B). Taken together, the disturbed Ca^2+^ kinetics in LVNC-CM, the significant Ca^2+^ leakage in DCM-CM and the diametrically opposed aberrations in systolic Ca^2+^ content indicates that different RBM20 mutations dictate different impairments in Ca^2+^ homeostasis.

Therefore, we next investigated the impact of the distinct RBM20 mutations on contractility and force generation. Engineered human myocardium (EHM) were produced from iPSC-CM and commercially available human foreskin fibroblasts (Fig. 4A). The EHM were cast as a 3D ring structure around two flexible poles and allowed to mature for 4-6 weeks. The isogenic resDCM1 line served as control. During the 4 weeks of EHM maturation, the beating frequency is optically captured and reaches a plateau after 3-4 weeks. At day 35, the LVNC and the DCM1-EHM exhibit a significantly faster beating rate than the corresponding rescue line (Fig. 4B). As a functional end-point assessment, the EHM were measured under isometric conditions in organ baths. Both LVNC-as well as the DCM1-EHM showed a reduced force of contraction (FOC) (active force) which was more pronounced in LVNC-EHM (Fig. 4C). The resting force of EHM (passive force) was significantly increased for DCM-EHM suggesting a higher stiffness (Fig. 4D). Importantly, the active/passive force ratio as an indicator of heart muscle dysfunction was reduced in both, LVNC- and DCM-EHM, compared to resDCM1-EHM underscoring a pathological contractility when an RBM20 mutation is present (Fig. 4E). Lastly, the contraction and relaxation time was accelerated for LVNC- and DCM-EHM with LVNC exhibiting the most prominent decrease (Fig. 4F/G), which is in line with the accelerated Ca^2+^ kinetics data.

**Fig. 4:**
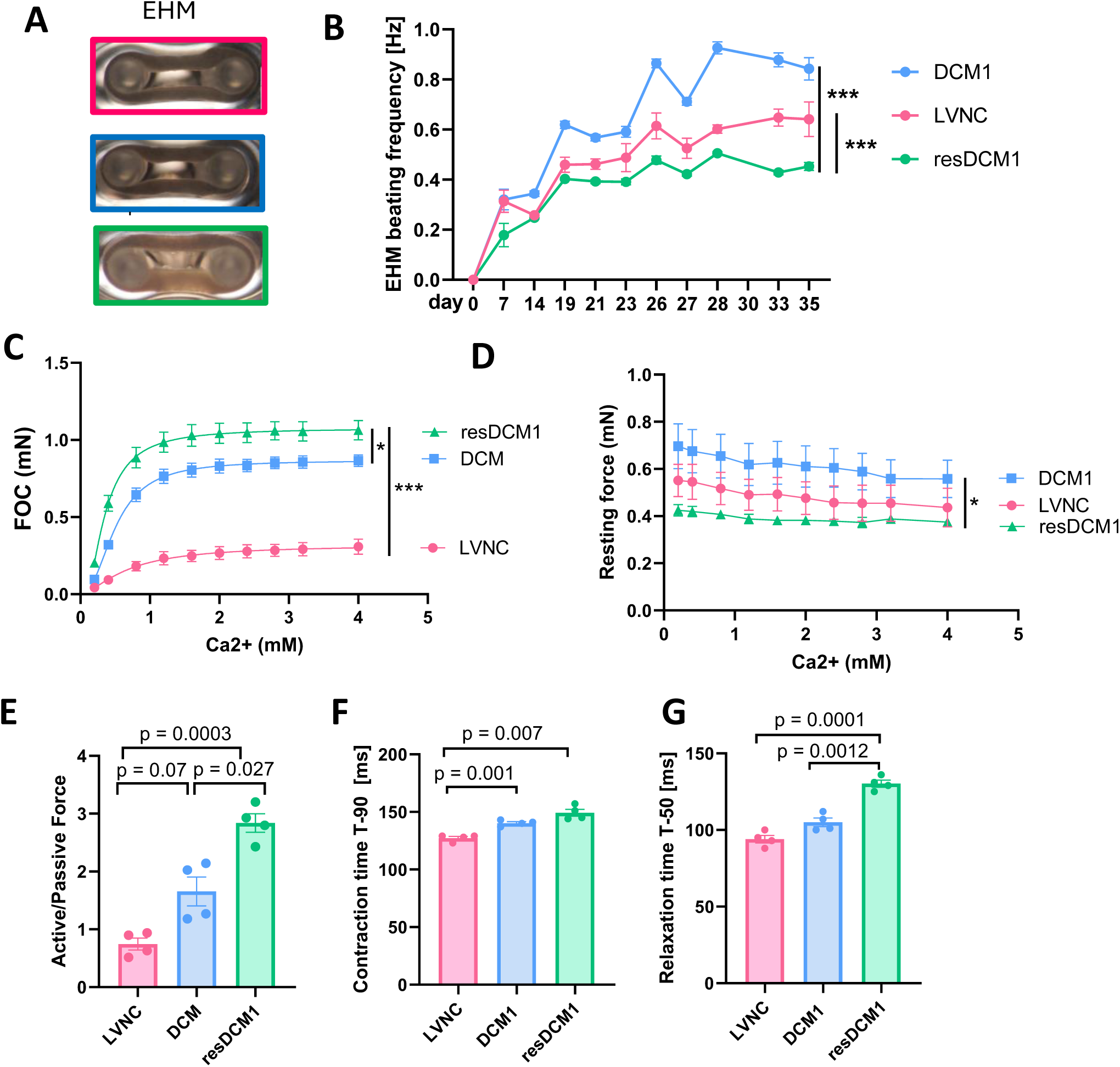
LVNC- and DCM-EHM show differential Ca^2+^ handling pathologies and contractile dysfunction. **A:** Engineered human myocardium (EHM) for LVNC, DCM1 and resDCM1 on flexible holders. **B:** Beating rate of the EHM me asured on different days since EHM casting (day 0 refers to the day of casting). 14-16 EHM per cell line. Mann-Whitney test for day 35 resDCM1 vs LVNC and resDCM1 vs DCM1. **C-E:** 4 EHM were used for isometric force measurements for each line. **C:** Force of contraction (FOC) measurements under rising Ca^2+^ concentrations. **D:** Resting Force measurements under rising Ca^2+^ concentration. **C and D:** P-values were calculated by Two-way RM ANOVA with Geisser-Greenhouse,s correction. P-value (column factor) against resDCM1 vs DCM1 and resDCM1 vs LVNC (significant values are marked with * p<0.05, ** p<0.01, ***p<0.001). **E-G:** P-value by Brown-Forsythe and Welch ANOVA with Dunnett,s multiple comparisons test. **E:** Active/passive force (at 4 mM Ca^2+^) calculation. **F:** Contraction time (T-90: to 90 % of baseline) of EHM in ms. **G:** Relaxation (T-50: to 50 % of baseline) time of EHM in ms.

### Increased cAMP levels and hyperphosphorylation events are specific to LVNC-CM, but not DCM-CM

Due to RBM20-dependent alterations in Ca^2+^ homeostasis, we sought to investigate the β-adrenergic signaling pathway including the second messenger cAMP as well as protein kinase A (PKA) and CAMK2D activity. CAMP dynamics were measured by transduction of iPSC-CM with an adenoviral construct expressing the Forster resonance energy transfer (FRET)-based cAMP sensor Epac1-camps under the control of the cytomegalovirus promoter [35]. First, we analyzed cAMP synthetization under basal conditions by pretreating iPSC-CM with the unselective phosphodiesterase (PDE) inhibitor 3-isobutyl-1-methylxanthin (IBMX) in the absence of Iso (Fig. 5A). We detected significantly higher FRET responses in LVNC-CM compared to DCM-CM that were rescued in resLVNC-CM (Fig. 5B) suggesting increased basal cytoplasmic cAMP levels in LVNC-CM. Upon Iso treatment, we did not observe differences in CFP/YFP ratios between the groups (Suppl. Fig. 6C). Next, we explored the phosphorylation status of proteins involved in Ca^2+^ cycling. One protein that heavily influences Ca^2+^ reuptake into the SR, and therefore the Ca^2+^ transient decay time, is the SERCA-inhibitory protein phospholamban (PLN). PLN,s inhibitory effect can be ameliorated by phosphorylation by protein kinase A (PKA) of serine 16 (S16p) and CAMK2D of threonine 17 (T17p). Therefore, we focused on the phosphorylation status of PLN in basal and Iso-treated samples. At baseline, the LVNC-CM exhibited higher phosphorylation of PLN-S16p by PKA and PLN-T17p by CAMK2D compared to resLVNC-CM although that tendency was not significant for the PKA site (p=0.057) (Fig. 5C/D/E). As expected, Iso-treated iPSC-CM showed a prominent phosphorylation increase in all samples (Fig. 5C). However, compared to the phosphorylation increase in resLVNC, the LVNC-CM showed a significantly lower rise in phosphorylation events, which could be due to higher phosphorylation levels at basal state, especially for the CAMK2D site (Suppl. Fig. 7A). In contrast, the increase in PLN-S16p phosphorylation after Iso stimulation is comparable between LVNC- and resLVNC-CM (Suppl. Fig. 7A), indicating that the CAMK2D-dependent PLN-T17p site is affected to a higher extent. Like the cAMP findings, DCM-CM do not show any deviations in phosphorylation status of PLN before and after Iso treatment (Fig. 5C/D/E, Suppl. Fig. 7B).

**Fig. 5:**
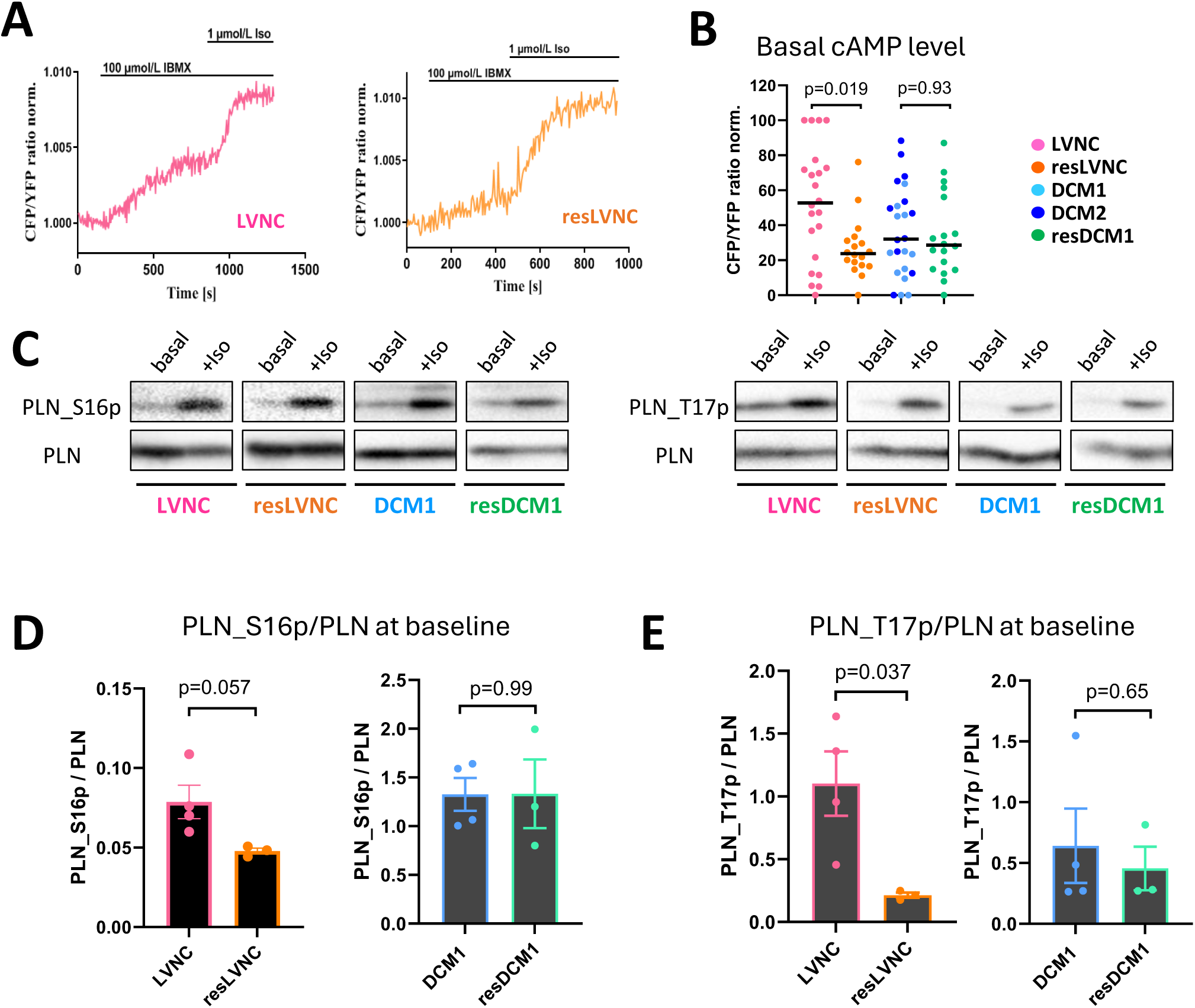
LVNC-CMs exhibit increased cAMP levels and altered PKA- and CAMK2D-dependent phosphorylation of PLN. **A:** Representative cAMP FRET traces from the EPAC-cAMP-FRET sensor in adenovirally transduced iPSC-CM from indicated patient groups at basal level. The maximal FRET response was induced using the unselective phosphodiesterase (PDE) inhibitor (IBMX) (100 µM). **B:** LVNC-CM show elevated basal cAMP levels. Quantification of maximal FRET response to IBMX at basal state. P-value was calculated by Mann-Whitney test patient vs rescue line. [number of differentiations/analyzed cell]: LVNC [4/22], resLVNC [3/18], DCM1+2 [4/24] and resDCM1 [3/18]. **C:** Representative original Western blot membranes stained for PLN and its PKA-phosphorylation site S16p (left) and its CAMK2D-phosphorylation site T17p (right). Samples were taken unstimulated (basal) and after Iso treatment (1 µM for 15 min). **D+E:** Quantification of PLN-S16p and PLN-T17p at baseline. Every dot represents one cardiac differentiation. The p-values were calculated by Students t-test.

In conclusion, under basal conditions, key regulatory enzymes involved in cytosolic Ca^2+^ handling are hyperphosphorylated in LVNC-CM, but not DCM-CM. In LVNC-CM, the elevated levels of cAMP with subsequently higher phosphorylation of PLN by PKA and CAMK2D could be the driver for the distinctly fastened Ca^2+^ kinetics in LVNC-CM.

### L-type Ca^2+^ current blocker verapamil in the therapy of RBM20 based cardiomyopathies

Based on the distorted Ca^2+^ handling, we explored possibilities for drug intervention. Based on their clinical phenotype of HFrEF, the current guidelines for both families would include β-blocker and additional heart failure medications including angiotensin receptor blocker (candesartan), the mineralocorticoid receptor antagonist eplerenone, but not calcium blockers (Suppl. Fig. 7C). Since Ca^2+^ handling was a prominent feature in these cardiomyopathies, we interrogated whether Ca^2+^ blockage could still be beneficial. We compared the effects of verapamil with the effects of the β-blocker (metoprolol), since cAMP and CAMK2D signaling – both under the control of β-adrenergic stimulation [36]– were also altered in both cell lines. In fact, treatment with verapamil (30 nM) accelerated Ca^2+^ rise kinetics and restored the physiological response to β-adrenergic stimulation in LVNC-CM (Fig. 3E). These effects were more pronounced for verapamil than for metoprolol (5 µM), as metoprolol treatment only restored the Iso response but did not affect basal Ca^2+^ kinetics (Fig. 3E). Furthermore, verapamil (30 nM) and metoprolol (5 µM) significantly decrease the Ca^2+^ leak in DCM-CM (Fig. 3F). In summary, verapamil is an interesting antiarrhythmic candidate for both RBM20-mutation-based pathologies regarding Ca^2+^ cycling and Ca^2+^ leakage.

### Increased mitochondrial respiration in LVNC-CM, but not DCM-CM

Besides genes involved in EC coupling, RBM20 also targets mitochondrial genes, and in LVNC-CM, *IMMT* was mis-spliced. Therefore, we analyzed the impact of both mutations on mitochondrial and metabolic function. Visualizing the mitochondrial network with Mitospy revealed that compared to control, this was more prominently developed in LVNC-CM, but not DCM-CM (Fig. 6A/B). Also, the mitochondrial membrane potential (ΔΨm; determined by TMRM) was increased in LVNC-CM, but not DCM-CM, compared to control (Fig. 6C). Accordingly, both basal and maximal oxygen consumption rate (OCR) were elevated in LVNC-CM, but not DCM-CM (Fig. 6D/E). All these effects were rescued in the resLVNC lines, confirming the specificity of the RBM20 mutation for these alterations. Since the elevated respiratory capacity in LVNC-CM may be related to the elevated energetic demand imposed by activation of EC coupling, we interrogated whether verapamil or metoprolol would also normalize elevated metabolic activity. In fact, both compounds normalized the elevated basal, but not the maximal respiratory capacity in LVNC-CM (Fig. 6F). These results suggest that elevated basal OCR in LVNC-CM could be, at least in part, a secondary effect due to enhanced cytosolic Ca^2+^ handling and not a direct cause of the RBM20 mutation per se.

**Fig. 6:**
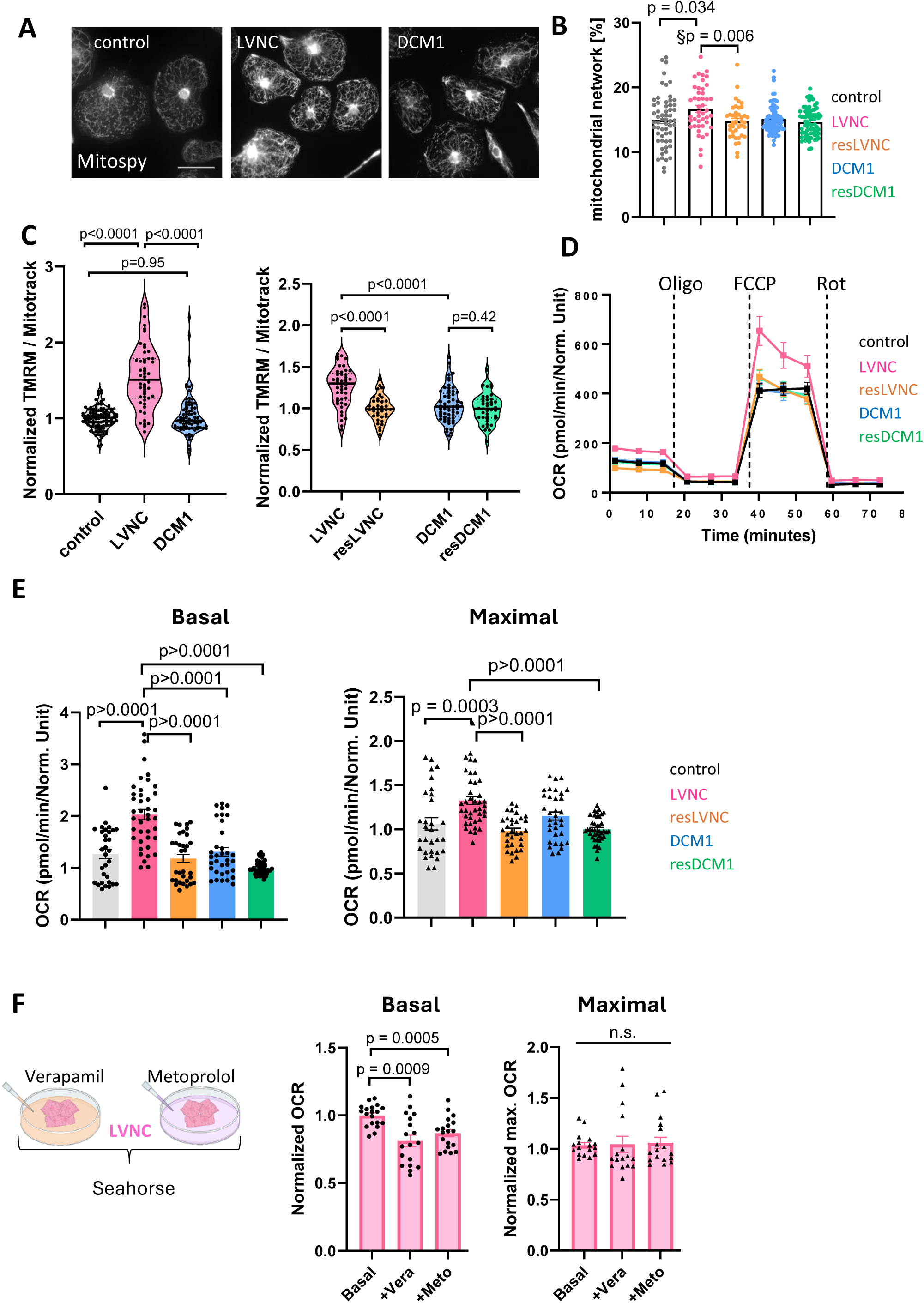
Mitochondrial and metabolic profiling of LVNC- and DCM-CM. **A:** Representative Mitospy-stainings. Brightness and contrast is enhanced for visualization purposes. Scale bar: 50 µm. **B:** Quantified mitochondrial network in [%] using the MiNa ImageJ plugin. [number of differentiations/analyzed images] for control [4/59], LVNC [3/45], resLVNC [3/40], DCM1 [4/81] and resDCM1 [4/71]. P-values were calculated by Kruskal-Wallis test with Dunn,s correction against control. §P-values were calculated by Mann-Whitney test LVNC vs resLVNC and DCM1 vs resDCM1. Only significant p-values are noted. **C:** TMRM measurement of mitochondrial membrane potential. To compare measurements from different days the samples were either normalized to control or to their respective isogenic line. LVNC-CM show increased mitochondrial membrane potential. [number of differentiations/analyzed images] for left: control [7/110], LVNC [3/51] and DCM1 [5/79]. Right: LVNC [2/45], resLVNC [2/36], DCM1 [3/60] and resDCM1 [2/40]. P-values were calculated by Mann-Whitney test. **D:** Representative Seahorse-curves. **E:** Quantification of basal and maximal respiration capacity for [number of differentiations/analyzed wells] control [4/31], LVNC [5/40], resLVNC [4/32], DCM1 [4/32] and resDCM1 [5/40]. To display different measurements in one graph, the data was normalized to resDCM1, which was part of every measurement. P-values were calculated by Kruskal-Wallis test with Dunn,s multiple comparisons test. **F:** LVNC-CM were treated with verapamil (Vera) (30 nM) or metoprolol (Meto) (5 µM). Data is normalized to the basal condition. [Number of differentiations/analyzed wells] for basal [3/19], Vera [3/18] and Meto [3/19]. N.s.: non significant; Vera: verapamil; Meto: metoprolol

### DCM mutation p.R634W in LVNC background mirrors the DCM phenotype

Since the two RBM20 variants show distinct phenotypes, we used the resLVNC-line to introduce the DCM-variant p.R634W using CRISPR/Cas9 (Suppl. Fig. 8A). Similar to all the other lines, the newly generated iPSC are positive for pluripotency markers (Suppl. Fig. 8B) and retained a normal karyotype (46, XX) after the second gene edit (Suppl. Fig. 8C). This novel line, termed resLVNC-DCM.W, represents the DCM variant but now in a LVNC background to address the question of whether the DCM variant per se will cause a DCM phenotype or if the genetic background of the LVNC patient contributes to the development of the LVNC phenotype. Intriguingly, the resLVNC-DCM.W-CM exhibited a phenotype closer to the DCM than the LVNC. We observed decreased sarcomeric regularity (Fig. 7A) and no alteration in ΔΨm (Fig. 7B). Furthermore, the force of contraction of the resLVNC-DCM.W-EHM was virtually identical to the force generation of the DCM-EHM, and resting force similarly elevated as in the DCM-EHM, but different from the strong hypo-contractile phenotype of the LVNC-EHM (Fig. 7C/D). Accordingly, also the active/passive force ratio and the contraction and relaxation time resembled the DCM-EHM rather than the LVNC-EHM phenotype (Fig. 7E/F/G).

**Fig. 7:**
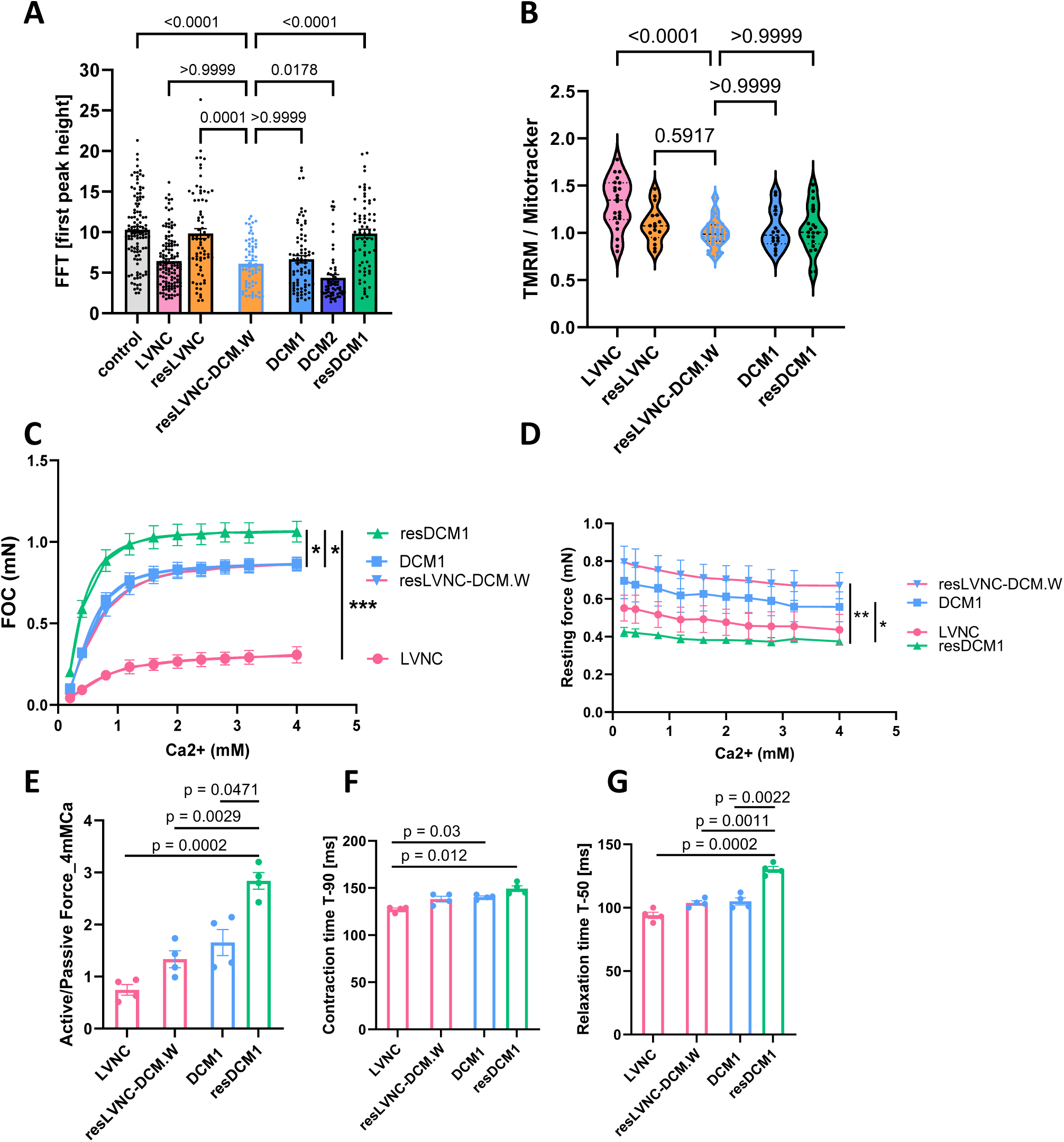
ResLVNC-DCM.W mirror the DCM phenotype. **A:** Quantification of Z-disc sarcomeric regularity (α-actintin) using Fast Fourier transformation. Bar graphs depict the peak amplitude of the first-order peak. Previous graph from Fig. 2E with new data for resLVNC-DCM.W. [number of differentiations/analyzed pictures] for control [6/112], LVNC [7/128], resLVNC [4/73], <u>resLVNC-DCM.W [3/66],</u> DCM1 [4/79], DCM2 [3/58] and resDCM1 [4/69]. Data is represented as bar graph with mean +/-SEM. P-values were determined by Kruskal-Wallis test against resLVNC-DCM.W with Dunn,s correction. **B:** TMRM measurement of mitochondrial membrane potential. To compare measurements from different days the samples were normalized to resLVNC-DCM.W. [number of differentiations/analyzed pictures] for LVNC [1/22], resLVNC [1/16], <u>resLVNC-DCM.W [2/43],</u> DCM1 [1/16] and resDCM1 [1/21]. Data is represented as violin plot. P-values were determined by Kruskal-Wallis test against resLVNC-DCM.W with Dunn,s correction. **C-E:** EHM data for resLVNC-DCM.W (n=4). ResLVNC-DCM.W resembles the DCM phenotype and differs from LVNC. **C:** Force of contraction (FOC) measurements under rising Ca^2+^ concentrations. The resLVNC-DCM.W-EHM is almost congruent with the DCM1-EHM measurements. P-values were calculated by Two-way RM ANOVA with Geisser-Greenhouse,s correction. P-value (column factor) for resDCM1 vs DCM1; LVNC vs resDCM1; resDCM1 vs resLVNC-DCM.W (significant values are marked * p<0.05, ** p<0.01, ***p<0.001) and DCM1 vs resLVNC-DCM.W (not significant). **D:** Resting Force measurements under rising Ca^2+^ concentration. P-values were calculated by Two-way RM ANOVA with Geisser-Greenhouse,s correction. P-value (column factor) for resDCM1 vs DCM1 and resDCM1 vs resLVNC-DCM.W (significant values are marked * p<0.05, ** p<0.01, ***p<0.001) and LVNC vs resDCM1 and DCM1 vs resLVNC-DCM.W (not significant). **E-G:** P-value by Brown-Forsythe and Welch ANOVA with Dunnett,s multiple comparisons test. Only significant values are noted. **E:** Active/passive force (at 4 mM Ca^2+^) calculation. **F:** Contraction time (T-90: to 90 % of baseline) of EHM in ms. **G:** Relaxation (T-50: to 50 % of baseline) time of EHM in ms.

## Discussion

In a human patient-specific approach, the present study shows that two distinct missense mutations in the same amino acid of RBM20 (R634) can lead to very different phenotypes of EC coupling and mitochondrial energetics. On the one hand, both mutations cause molecular and cellular deviations such as the aberrant cytoplasmatic accumulation of RBM20, erroneous splicing of *TTN* and *RYR2,* and sarcomeric disarray. In contrast, depending on the respective nucleic acid substitution (Trp or Leu), the ensuing phenotype in EC coupling and metabolism was very different. The DCM-CM with the Trp substitution generated a phenotype that was rather reminiscent of the “classical” EC coupling phenotype as in HFrEF, with decreased peak systolic Ca^2+^ concentrations, an increased SR Ca^2+^ leak and modest reduction in systolic force generation but increased diastolic force in a 3D EHM model. In this DCM model, metabolic parameters were largely unchanged. Conversely, in the LVNC mutation, peak systolic Ca^2+^ was increased, together with accelerated kinetics of Ca^2+^ rise and decay, mediated by increased baseline cAMP levels and PKA- and CAMK2D-dependent phosphorylation of PLN. In parallel to this adrenergic pre-activation, also basal and maximal respiratory capacity, mitochondrial membrane potential, and network density were substantially enhanced in LVNC-CM. Intriguingly, despite this activation of Ca^2+^ handling, systolic force generation was strongly reduced. These data indicate that the relevant contractile deficit observed in the 3D EHM is likely related to factors downstream of cytosolic Ca^2+^ handling. Future studies must clarify whether this systolic deficit results from alterations of sarcomere structure or function.

A significant debate in RBM20 pathology is whether the RBM20-dependent erroneous splicing, the RBM20 cytoplasmic accumulation, or the RBM20 abundance is the stronger or predominant disease driver. For all RBM20 mutations and knock-out models, mis-splicing was shown for multiple target genes that always included *TTN* [10]. On the other hand, recent reports highlighted that aberrant cytoplasmatic RBM20-granulae formation is the pathology’s culprit, whereby restoring nuclear localization of mutant RBM20 rescued the mis-splicing and RBM20 phenotype [37]. However, for RBM20 variants outside the RS domain mis-splicing without RBM20 cytoplasmic accumulation was shown, which also resulted in DCM phenotype [10]. Here, we report RBM20-granulae for both missense variants and shared mis-splicing for a number of target genes exemplified by *TTN* and *RYR2*. Additionally, distinct mis-splicing events occur for other targets depending on the RBM20 variant, e.g. for Ca^2+^ handling genes (*CAMK2D*, *TRDN*) in LVNC-CM concomitant with a distinct Ca^2+^ handling phenotype. This indicates that, at least for the differential pathologies observed for these two variants, the differential mis-splicing is a stronger driver since the cytoplasmatic RBM20 aggregation was observed for both variants. This should be taken with caution as the mis-localization and RBM20-granulae formation require deeper investigation regarding its spatiotemporal resolution and organelle interaction. Aberrant granulae formation underlies many neurodegenerative diseases, in which temporal and dynamic localization of aggregates are key to understanding the pathomechanism [38, 39]. As a common theme, we see *TTN* and sarcomeric disarray, which is a shared pathology for all RBM20 variants [10]. The sequencing data support this hypothesis as overall gene expression profiles do not substantially differ between LVNC- and DCM-specific phenotypes in Ca^2+^ and metabolic homeostasis. Rather the differential exon usage of RBM20 target genes shows more pleiotropic differences between RBM20-mutation dependent LVNC and DCM.

In addition to sarcomeric disarray, Ca^2+^ handling aberrations are common in RBM20-dependent cardiomyopathies. Ca^2+^ analysis revealed distinct pathological phenotypes for different RBM20 mutations, which were reversed in isogenic lines, highlighting that RBM20 mutations can lead to varied outcomes. Systolic Ca^2+^ was elevated for LVNC-CM and adversely significantly reduced for DCM-CM. A study in RBM20 knock-out mouse models reported increased systolic Ca^2+^, SR Ca^2+^-load, and Ca^2+^ leakage [19] suggesting that full RBM20 knockout affects multiple aspects of Ca^2+^ homeostasis, while single mutations do not. For example, LVNC-CM showed elevated systolic Ca^2+^ but no Ca^2+^ leakage, while DCM-CM showed Ca^2+^ leakage but reduced systolic Ca2+. Our former study in iPSC-CM with RBM20-p.S635A reported decreased diastolic Ca^2+^ and increased systolic Ca^2+^ levels [17], which is reminiscent of the LVNC-CM described here. However, iPSC-CMs with p.S635A and p.P633L reported increased Ca^2+^ transient rise times [17, 24], whereas the LVNC-CM adversely presented with distinctly fastened/decreased Ca^2+^ cycling times. In contrast to sarcomeric disarray, which is uniformly reported in RBM20-dependent cardiomyopathy, Ca^2+^ handling parameters show differential pathologies in LVNC and DCM-CM, which depend on the underlying missense mutations. This is the first report that shows the distinct pathologies in a direct side-by-side comparison. The Ca^2+^ data portray distinct aberrations for LVNC and DCM, while LVNC-CM shows signs of hypercontractility with fast cycling times and elevated systolic Ca^2+^, the DCM-CM align more with a heart failure phenotype (decreased systolic Ca^2+^ and Ca^2+^ leak). Here we expand this pathology to an overstimulated B-adrenergic system in LVNC-CM with elevated cAMP levels. The sequencing data revealed that enzymes involved in cAMP generation are affected by the RBM20 mutations, especially adenylate cyclase 6 (data not shown). However, there was no predicted difference between LVNC and DCM. Therefore, this does not explain the distinctly elevated cAMP levels in LVNC-CM.

The Ca^2+^ channel blocker Verapamil showed promising potential by positively influencing multiple aspects of Ca^2+^ cycling and Ca^2+^ leakage. This is in line with the previous observation that verapamil treatment attenuated Iso-induced Ca^2+^ leakage in RBM20 knockout mice [19]. One major caveat is that Verapamil is contraindicated for patients with HFrEF since it has profound negative inotropic effects. Still, it might be a valuable candidate to explore its application in pre-heart failure RBM20 patients. This mandates further studies and randomized clinical investigations. Another study already provided evidence that drug testing in patient-specific iPSC-CM and a tailored medication switch is beneficial to the patient [18]. This underscores how iPSC-CM represent a great tool in cardiac personalized medicine. The EHM results show uniformly decreased FOC for LVNC- and DCM-CM, with LVNC-CM showing the poorest performance. The EHM data is suggestive of two hypotheses: First, Ca^2+^ content correlates with force generation, but although the systolic Ca^2+^ contents differed significantly for LVNC and DCM, the FOC is decreased for both, which indicates that the deranged sarcomeric structure plays a key role in failing to produce sufficient force. Second, although the end result of reduced heart performance is the same for the patients, the road to this may be different due to different underlying calcium deviations. In conclusion, both RBM20 mutations cumulate in significantly impaired force generation, which is in line with the patient,s pathology of severely reduced EF. Further, it should be noted that rescue of the gene mutation reversed all phenotypes observed, suggesting that gene and base editing, especially for RBM20, is a feasible way forward. Although this is not available in the clinics yet, the first reports show promising results [40].

In contrast to distinct Ca^2+^ phenotypes, enhanced metabolic activity was uniquely observed for LVNC-CM. This is the first study that reports metabolic aberrations for a RBM20 missense variant and concomitantly shows that this impairment is not a general phenotype for all RBM20 variants. A recent study shows that metabolic activity declines for the rare loss-of-function RBM20 mutations [22]. However, treatment with verapamil or metoprolol attenuated the enhanced basal OCR but did not affect the maximal OCR. This suggests that the metabolic phenotype is, at least in part, influenced by the calcium aberrations, although it cannot be completely excluded that these two drugs have also direct effects on the metabolism as well. Our preliminary data show that mitochondrial calcium may be influenced since mitochondrial Ca^2+^ importer subunits of the mitochondrial Ca^2+^ uniporter (MCU), as well as the mitochondrial Ca^2+^ exporter NCLX, are downregulated in expression in LVNC-CM (data not shown). Since it has been shown how granulae can interact and damage mitochondria in neurodegenerative diseases [39], the question arises whether this is also a mechanism in cardiac diseases. Resolving these granulae or investigating RBM20 variants without mis-localization regarding their metabolism should be addressed in future studies. In conclusion, similar to Ca^2+^ handling aberrations, metabolic phenotypes depend on the RBM20 variant, and therefore, not all mutation carriers would, for example, benefit from medication that targets metabolism.

**In conclusion, RBM20 variants are often discussed as one entity. Still, this report highlights that mutation matters and results in different pathologies, which should be considered when moving forward in personalized medicine.**

### Study limitations

A limitation of the current study is the inherent immaturity of iPSC-CM. It has been shown that iPSC-CM resemble fetal-like CM more closely than adult CM and could therefore mask possible phenotypes [41]. In this study, we use prolonged culture times of 60-90 days or culture in 3D EHM constructs to enhance maturation, as previously shown [42, 43]. However, full CM maturity has not been achieved yet, and new strategies to enhance maturation further need to be considered. Furthermore, the patient’s genetic background, sex and age may have an influence, which we circumvented in part by using isogenic lines in addition to control lines.

## Supporting information

Supplements Rebs et al

## Acknowledgments

The authors thank Johanna Heine, Sandra Georgi, and Yvonne Metz for their excellent technical assistance, as well as anybody who helped take skin punch biopsies from the DCM patient. This work was supported by the Bundesministerium fur Bildung und Forschung (BMBF) grant [CaRNAtion 031L0075C to KSB, GH, BM, KG], the German Center for Cardiovascular Research (DZHK) [to KSB and BM], the German Heart Foundation/German Foundation of Heart Research [AZ. F/38/18] (to KSB), the Else Kroner-Fresenius-Stiftung Foundation [2017-A137] (to KSB), the Fritz Thyssen Foundation Az 10.19.2.026MN to KSB and the Deutsche Forschungsgemeinschaft (to KSB). CM is supported by the DFG (SFB 1525, project #453989101; Ma 2528/8-1, project #505805397). MS received funding from the European Research Council (ERC) under the European Union’s Horizon 2020 research and innovation program (grant agreement No 835102). SR received funding from the German Cardiac Society (DGK) (Project-No DGK07/2024).

## Author contributions

SR generated and characterized the iPSC and CRISPR/Cas9 lines performed most experiments and drafted and wrote the manuscript. FS, EK, and BM provided patient material and patient data. HE generated and characterized the CRISPR-edit line. VON provided the FRET sensors and DH generated the FRET data. CR and BM generated and analyzed the RNAseq data. MS, SD and TK generated and analyzed the dSTORM data. CM and JD performed and analyzed the Seahorse data. BM received funding and developed the concept. KG and GH contributed to study design and received funding. KSB developed the concept, designed the study, received funding, and wrote the manuscript.

## References

1. McNally, E.M. and L. Mestroni, Dilated Cardiomyopathy: Genetic Determinants and Mechanisms. Circ Res, 2017. 121(7): p. 731–748.

2. Arbelo, E., et al., 2023 ESC Guidelines for the management of cardiomyopathies. Eur Heart J, 2023. 44(37): p. 3503–3626.

3. Kayvanpour, E., et al., *Clinical and genetic insights into non-compaction: a meta-analysis and systematic review on* 7598 *individuals*. Clin Res Cardiol, 2019. 108(11): p. 1297–1308.

4. Wilcox, J.E. and R.E. Hershberger, Genetic cardiomyopathies. Curr Opin Cardiol, 2018. 33(3): p. 354–362.

5. Meyer, H.V., et al., Genetic and functional insights into the fractal structure of the heart. Nature, 2020. 584(7822): p. 589–594.

6. Sedaghat-Hamedani, F., et al., Clinical genetics and outcome of left ventricular non-compaction cardiomyopathy. Eur Heart J, 2017. 38(46): p. 3449–3460.

7. Jordan, E., et al., Evidence-Based Assessment of Genes in Dilated Cardiomyopathy. Circulation, 2021. 144(1): p. 7–19.

8. Li, D., et al., Identification of novel mutations in RBM20 in patients with dilated cardiomyopathy. Clin Transl Sci, 2010. 3(3): p. 90–7.

9. Parikh, V.N., et al., Regional Variation in RBM20 Causes a Highly Penetrant Arrhythmogenic Cardiomyopathy. Circ Heart Fail, 2019. 12(3): p. e005371.

10. Gregorich, Z.R., et al., Mechanisms of RBM20 Cardiomyopathy: Insights From Model Systems. Circ Genom Precis Med, 2024. 17(1): p. e004355.

11. Dai, J., et al., RBM20 Is a Candidate Gene for Hypertrophic Cardiomyopathy. Can J Cardiol, 2021. 37(11): p. 1751–1759.

12. Inagaki, N., et al., Pathogenic variant of RBM20 in a multiplex family with hypertrophic cardiomyopathy. Hum Genome Var, 2022. 9(1): p. 6.

13. Beqqali, A., et al., A mutation in the glutamate-rich region of RNA-binding motif protein 20 causes dilated cardiomyopathy through missplicing of titin and impaired Frank-Starling mechanism. Cardiovasc Res, 2016. 112(1): p. 452–63.

14. Takahashi, K., et al., Induction of pluripotent stem cells from adult human fibroblasts by defined factors. Cell, 2007. 131(5): p. 861–72.

15. Gahwiler, E.K.N., et al., Human iPSCs and Genome Editing Technologies for Precision Cardiovascular Tissue Engineering. Front Cell Dev Biol, 2021. 9: p. 639699.

16. Borchert, T., et al., Catecholamine-Dependent beta-Adrenergic Signaling in a Pluripotent Stem Cell Model of Takotsubo Cardiomyopathy. J Am Coll Cardiol, 2017. 70(8): p. 975–991.

17. Streckfuss-Bomeke, K., et al., Severe DCM phenotype of patient harboring RBM20 mutation S635A can be modeled by patient-specific induced pluripotent stem cell-derived cardiomyocytes. J Mol Cell Cardiol, 2017. 113: p. 9–21.

18. Prondzynski, M., et al., Disease modeling of a mutation in alpha-actinin 2 guides clinical therapy in hypertrophic cardiomyopathy. EMBO Mol Med, 2022. 14(8): p. e16423.

19. van den Hoogenhof, M.M.G., et al., RBM20 Mutations Induce an Arrhythmogenic Dilated Cardiomyopathy Related to Disturbed Calcium Handling. Circulation, 2018. 138(13): p. 1330–1342.

20. Guo, W., et al., *RBM20*, *a gene for hereditary cardiomyopathy*, *regulates titin splicing*. Nat Med, 2012. 18(5): p. 766–73.

21. Schneider, J.W., et al., Dysregulated ribonucleoprotein granules promote cardiomyopathy in RBM20 gene-edited pigs. Nat Med, 2020. 26(11): p. 1788–1800.

22. Vad, O.B., et al., Loss of Cardiac Splicing Regulator RBM20 Is Associated With Early-Onset Atrial Fibrillation. JACC Basic Transl Sci, 2024. 9(2): p. 163–180.

23. Wyles, S.P., et al., Modeling structural and functional deficiencies of RBM20 familial dilated cardiomyopathy using human induced pluripotent stem cells. Hum Mol Genet, 2016. 25(2): p. 254–65.

24. Briganti, F., et al., iPSC Modeling of RBM20-Deficient DCM Identifies Upregulation of RBM20 as a Therapeutic Strategy. Cell Rep, 2020. 32(10): p. 108117.

25. Fenix, A.M., et al., Gain-of-function cardiomyopathic mutations in RBM20 rewire splicing regulation and re-distribute ribonucleoprotein granules within processing bodies. Nat Commun, 2021. 12(1): p. 6324.

26. Rebs, S., et al., Generation of pluripotent stem cell lines and CRISPR/Cas9 modified isogenic controls from a patient with dilated cardiomyopathy harboring a RBM20 p.R634W mutation. Stem Cell Res, 2020. 47: p. 101901.

27. Heilemann, M., et al., Subdiffraction-resolution fluorescence imaging with conventional fluorescent probes. Angew Chem Int Ed Engl, 2008. 47(33): p. 6172–6.

28. van de Linde, S., et al., Direct stochastic optical reconstruction microscopy with standard fluorescent probes. Nat Protoc, 2011. 6(7): p. 991–1009.

29. Doose, S., LOCAN: a python library for analyzing single-molecule localization microscopy data. Bioinformatics, 2022. 38(9): p. 2670–2672.

30. Schubert, E., et al., *DBSCAN Revisited*, *Revisited*. ACM Transactions on Database Systems, 2017. 42(3): p. 1–21.

31. Rebs, S., T.A. Buchwald, and K. Streckfuss-Bomeke, A quantitative RT-PCR protocol to adapt and quantify RBM20-dependent exon splicing of targets at the human locus. STAR Protoc, 2022. 3(1): p. 101117.

32. Clark, K.A., et al., Striated muscle cytoarchitecture: an intricate web of form and function. Annu Rev Cell Dev Biol, 2002. 18: p. 637–706.

33. Bers, D.M., Cardiac excitation-contraction coupling. Nature, 2002. 415(6868): p. 198–205.

34. Eisner, D.A., et al., The Control of Diastolic Calcium in the Heart: Basic Mechanisms and Functional Implications. Circ Res, 2020. 126(3): p. 395–412.

35. Borner, S., et al., FRET measurements of intracellular cAMP concentrations and cAMP analog permeability in intact cells. Nat Protoc, 2011. 6(4): p. 427–38.

36. Grimm, M. and J.H. Brown, Beta-adrenergic receptor signaling in the heart: role of CaMKII. J Mol Cell Cardiol, 2010. 48(2): p. 322–30.

37. Kornienko, J., et al., Mislocalization of pathogenic RBM20 variants in dilated cardiomyopathy is caused by loss-of-interaction with Transportin-3. Nat Commun, 2023. 14(1): p. 4312.

38. Lim, J. and Z. Yue, *Neuronal aggregates: formation*, *clearance*, *and spreading*. Dev Cell, 2015. 32(4): p. 491–501.

39. Lurette, O., et al., Aggregation of alpha-synuclein disrupts mitochondrial metabolism and induce mitophagy via cardiolipin externalization. Cell Death Dis, 2023. 14(11): p. 729.

40. Grosch, M., et al., Striated muscle-specific base editing enables correction of mutations causing dilated cardiomyopathy. Nat Commun, 2023. 14(1): p. 3714.

41. Hong, Y., et al., Engineering the maturation of stem cell-derived cardiomyocytes. Front Bioeng Biotechnol, 2023. 11: p. 1155052.

42. Tiburcy, M., et al., Defined Engineered Human Myocardium With Advanced Maturation for Applications in Heart Failure Modeling and Repair. Circulation, 2017. 135(19): p. 1832–1847.

43. Emanuelli, G., et al., A roadmap for the characterization of energy metabolism in human cardiomyocytes derived from induced pluripotent stem cells. J Mol Cell Cardiol, 2022. 164: p. 136–147.

