## Supplements Rebs et al for "RBM20-variants induce distinct calcium handling and metabolic phenotypes in patient-specific stem cell models of dilated and non-compaction cardiomyopathy"

### Supplemental Materials

Methods
Supplemental tables
Supplemental figures

#### Methods

##### Ethical statement

For reprogramming into iPSC, human cardiac tissue or blood samples were obtained in compliance with the ethical committee of the University Medical Center Goettingen (Az -10/9/15) and the ethical committee of University Heidelberg (Ethical approval number: S-329/2012).

##### Cell culture

***Reprogramming***

The iPSC-lines for DCM1, resDCM1 [1] and healthy control cells [2] have previously been published by our group. For LVNC and DCM2, skin samples were obtained from the patients and reprogrammed with plasmids for LVNC or with Sendai virus for DCM2 (Suppl. Tab. 1). Details for reprogramming with plasmids and Sendai virus are published elsewhere [2].

***iPSC culture***

The iPSCs were cultured in monolayer in cell culture dishes coated for 30 min at 37 °C with Geltrex (167 mg/L) (Thermo Fisher Scientific). The medium Essential 8 (E8) (Thermo Fisher Scientific) was refreshed daily. Upon reaching 80-90 % confluency, the cells were passaged using 3-5 min Versene solution at 37 °C. After passaging, iPSC were cultured for 24 h in E8 medium supplemented with 2 µM (M = mol/l) Thiazovivin (TZV).

***iPSC-CM differentiation***

For iPSC-CM differentiation, iPSC were plated and subsequently cultured until they reached 80-90 % confluent monolayer, and Wnt signaling was modulated as previously described [1, 2]. Briefly, on day 0, the medium was exchanged for Differentiation medium (RPMI 1640 Glutamax (Thermo Fisher scientific) with 500 mg/L Albumin and 200 mg/L-ascorbic acid ) with the addition of 4 µM CHIR 99021 (Merck). On day 2, the medium was exchanged to Differentiation medium supplemented with 2 µM Wnt inhibitor IWP2 (Merck). On day 6, the medium was changed to Differentiation medium without any additions. On day 8, the medium was changed to Cardio Culture medium (RPMI-Glutamax + B27 supplement (Gibco, 17504044)) and was subsequently refreshed every other day. CM beating was observed around day 11. Between day 14-21, cells were replated in Cardio Digest medium (Cardio Culture medium with 10 % FCS and 2 µM TZV) onto Geltrex-coated 6 well plates at a cell number of 800.000 and after 24-48 h medium was replaced to glucose-deprived Selection medium (RPMI without HEPES and glucose (Thermo Fisher Scientific) with 4 mM lactate/HEPES, 500 mg/L Albumin and 200 mg/L L-Ascorbic acid) for 3-5 days. After the metabolic selection step, Cardio Culture medium was renewed twice a week. IPSC-CM were cultured for at least 60 days before used in experiments.

##### CRISPR/Cas9 editing of RBM20

The CRISPR/Cas9 gene editing using a crRNA-tracrRNA-Cas9 complex, HDR-template, ssDNA template and electroporation enhancer (all from IDT) are described in detail for resDCM1 [1]. The edit for resLVNC was performed identically with the crRNA: 5´-CTCACCGGACTACGAGA CAG-3´ and the HDR-template: 5´GGTGTGAAGATTCTAAATCCTGCTCCTTGGCTCCC TCACAGATATGGCCCAGAAAGGCCGCGGTCTCGTAGTCCCGTCAGCCGGTCACTCTCCCCGAGGTC CCACACTCCCAGCTTCACCTCC-3´. Gene edit was performed on LVNC-iPSC at a passage <30 . The editing efficiency was 45 % (32 positive clones of 72). The edit for resLVNC-634.W was performed with the crRNA: 5'-CGGTCTCGTAGTCCGGTGAG-3' and the HDR-template: 5'-GGTGTGAAGATTCTAAATCCTGCTCCTTGGCTCCCTCACAGATATGGCCCAGAAAGGCCGTGGTCTCG TAGTCCGGTGAGCCGGTCACTCTCCCCGGTCCCACACTCCCAGCTTCACCT-3'. The editing efficiency was 23.5 % (17 positive clones of 30).

#### Engineered Heart Muscle (EHM)

##### *EHM generation*

EHM was generated in accordance with previously published protocols with minor modifications [3, 4]. Briefly, iPSC-CM of high purity were combined with human foreskin fibroblasts (HFF-1, ATCC, SCRC-1041) in a 70:30 % ratio and resuspended in Iscove's medium supplemented with 4 % B27 minus insulin, MEM non-essential amino acids, 300 µM L-ascorbic acid 2-phosphate, 100 ng/mL IGF-1, 10 ng/mL FGF-2, 5 ng/mL $VEGF_{165}$. To prepare the collagen mixture required for tissue hydrogel formation, equal volumes of collagen (Collagen Solutions, FS22024) and RPMI 2x (Thermo Fisher Scientific, 51800) were mixed and neutralized with 0.1 N NaOH (Merck, 137031). The cell suspension was resuspended in an appropriate volume of the collagen mixture, with 180 µL of the subsequent hydrogel being gently pipetted in a ring-shape in each well of a custom 48 multi-well plate (myriamed GmbH, myrPlate Uniform, TM5-MED). For the first three days of tissue culture, the medium was supplemented with 5 ng/mL TGF-β1 (Peprotech, AF-100-21C) and exchanged daily. For the following 11 days the EHM medium was exchanged daily, which was subsequently exchanged for an adapted EHM maturation medium (MEM α, Thermo Fisher Scientific, 22561021, 4 % B27 minus insulin, MEM non-essential amino acids, 100 ng/mL IGF-1, 10 ng/mL FGF-2, 5 ng/mL $VEGF_{165}$) for the remainder of the tissue culture period. To monitor contractile activity by optical measurements of pole deflection, non-destructive imaging of the EHM plates was performed with a myrImager (myriamed GmbH).

##### *Isometric force measurements*

Measurements of contractile function were performed under isometric conditions in organ baths at 37 °C in gassed Tyrode´s solution (120 NaCl, 1 $MgCl_2$, 0.2 $CaCl_2$, 5.4 KCl, 22.6 $NaHCO_3$, 4.2 $NaH_2PO_4$, 5.5 glucose and 5.5 L-ascorbic acid; all in mM). EHM were electrically stimulated at 1.5 Hz with 5 ms square pulses of 150 mA and mechanically stretched at 62.5 µm intervals until reaching the maximum force amplitude. Responses to increasing extracellular calcium concentrations (0.2 to 4.0 mM) were investigated for each individual EHM. Ratios of active and passive force were obtained by comparing actively generated force with the resting tension of the EHM at saturating calcium levels.

#### Cell staining

##### *Fixation of the cells for IF and FLOW*

IPSC or iPSC-CM were plated onto Geltrex-coated coverslips or 8-chambered coverslip system (Cellvis). For fixation, the cells were washed once with 1xPBS and incubated 20 min with 4 % Histofix solution (Roth). After Histofix was discarded, the cells were washed with 1x PBS and 1-2 mL Blocking solution (1 % BSA in 1x PBS) was added. The cells were stored in Blocking solution at 4 °C until usage (up to 2 months).

For FLOW, one well with iPSC-CM was detached using 0.25 % trypsin/EDTA for 10 min at 37 °C, stopped with FCS (1:1) and transferred to a FCS centrifugation tube (all centrifugation steps at 200 g for 5 min). Cell pellets were washed once in 1x PBS and subsequently stored in Blocking buffer at 4 °C for up to one month.

***Antibody stainings for IF and FLOW***

For IF stainings, primary and secondary antibodies were diluted in Staining solution (0.1 % triton-x in 1 % BSA/PBS). Dilutions and all antibodies used are listed in suppl. Tab. 2. The secondary antibody dilutions ranged between 1:500 to 1:1000.

For FLOW- cTNT staining, the cells were incubated over night at 4 °C with the primary antibody cTNT in a 1:500 dilution in FACS buffer (1 % BSA/PBS with 0.1 % Triton-X). The next day, the cells were washed thrice with FACS buffer and subsequently incubated with the secondary antibody in a 1:1000 dilution in FACS buffer (Suppl. Tab. 2). The cells were washed thrice again in FACS buffer und diluted in 200 µL 1x PBS for measurements using a FACS Canto II with the following settings: 10000 events, forward scatter 228 V, side scatter 440 V, 488 Alexa Fluor laser 390 V. The detection threshold was set according to the blank sample and cTNT positive cells were presented as cTNT positive cells in [%].

#### Gene analysis

***RNA Isolation and cDNA synthesis***

Total RNA was isolated as described in the SV Total RNA Isolation System (Promega) according to the manufacturer's instructions for "RNA Isolation from fibrous tissue". The DNase-digestion step was expanded to 30 min at RT. 100-300 ng of RNA was used in first-strand cDNA synthesis using the iScript cDNA Synthesis Kit (BioRad).

***Quantitative PCR***

Quantitative PCR was performed using 1x iQ™ SYBR® Green Supermix (Bio-Rad) with 5 ng cDNA and 500 nM primers (each forward and reverse) and run on the CFX real-time PCR detection system (Bio-Rad). Primers used are listed in suppl. Tab. 3.

#### Sequencing

***Sanger Sequencing***

To sequence the RBM20-exon 9 locus, gDNA was isolated using the QIAamp DNA Mini Kit (Qiagen) following the manufacturer´s instructions. The gDNA was measured using a Nanodrop and adjusted to a value between 100 – 200 ng/µL. To amplify the exon 9 locus, a PCR was conducted with 2 µL cDNA in a PCR mix consisting of 1 µL forward and reverse primer (10 µM, suppl. Tab. 3), 5 µL Green buffer (5x) (Promega), 1.6 µL dNTP mix (10 mM) (Bioline), 0.1 µL Tag-polymerase (Promega) and 14.3 µL water (end volume 25 µL). The PCR was rum with 56 °C annealing temperature with 30 cycles. The PCR product was sent for Sanger sequencing to Mircosynth Seqlab (Göttingen).

***Illumina Sequencing***

RNA sequencing using poly(A) enrichment of the mRNA was performed for a total of 16 samples, comprising 3 LVNC, 5 resLVNC, 3 DCM1, and 5 resDCM1 samples. A median read depth of 76.52 million was achieved per pair-ended sequencing data, with a read length of 75 base pair. The reads were subjected to a quality check using the tool fastp [5, 6] The reads were then aligned to human genome GRCh38 using the tool Subread [7, 8] using the splice-aware option. BAM files were then used as an input to the *featureCounts* tool of the Subread package, to count the number of reads aligning to the gene features as defined in NCBI RefSeq annotation for hg38 (build 38.2). For the counting, only uniquely mapping fragments were considered. Read counts were imported in R package *DESeq2* [9] for normalization of sequencing depth and analysis of the differentially expressed genes between groups. Wald tests were used to calculate log2 fold changes and associated p-values, which were adjusted for multiple testing using the Benjamini-Hochberg procedure. Differentially expressed genes with an adjusted p-value $< 0.05$ were considered significant. DEXSeq was used for the analysis of differential exon usage, focusing on variations in exon-level expression patterns between groups. The exon counts were normalized, and generalized linear models (GLMs) were employed to test for significant differences in exon usage across conditions [10]. To account for hidden batch effects, Surrogate Variable Analysis (SVA) was performed. The normalized counts for a total of 28,395 genes are visualized as a volcano plot. Gene Ontology (GO) term enrichment analysis was performed on the differentially expressed genes identified through DESeq2 using Cytoscape with ClueGo plugin (settings: Ontologie KEGG, pathway with p-value $< 0.05$).

#### Karyotyping

Karyotyping was performed at Life & Brain (Bonn, Germany) via array-based genome-wide genotyping utilizing the Illumina BeadArray. Data was analyzed in GenomeStudio v2.0 (Illumina) with the cnvPartition 3.2.0 plug-in.

#### Western Blot

***Protein lysis***

Cell pellets were lysed in lysis buffer (20 mM Tris-HCl [pH 7.4], 200 mM NaCl, 20 mM NaF, 1 % Igepal, 1mM $Na_3VO_4$, 1mM DTT, 1x PhosStop, 1x cOmplete, EDTA-free protease inhibitor). The solution was incubated 30 min on ice, and subsequently centrifuged and the supernatant collected. Protein concentration was determined with the Pierce BCA protein assay kit according to manufactrurer` s instructions.

***SDS-electrophoresis, blotting and imaging***

20 µg protein were denaturated in Laemmli buffer at 37 °C for 30 min and separated in SDS-polyacrylamide gels (4-15 % Mini-Protean precast Gel; BioRad) with constant 15-20 mA/gel. Next, the proteins were transferred to either 0.45 µm nitrocellulose membrane (RBM20) or methanol activated 0.45 µm PVDF (PLN, PLN-S16p, PLN-T17p) membranes using a semi-dry transfer system (BioRad). Membranes were blocked in 5 % milk or 5 % BSA depending on the antibody (Suppl. Tab. 2). Primary antibodies were diluted in 1 % milk or 1 % BSA (Suppl. Tab. 2) and incubated over night at 4 °C. HRP-coupled secondary antibodies (Suppl. Tab. 2) were incubated for 1 h at room temperature. For imaging with the ChemiDoc XRS+ imaging system (BioRad), the membranes were incubated for 5 min with Immobiolon chemiluminescent HRP substrate (Millipore) and imaged with increasing exposure time.

#### Microscopy

##### *Brightfield*

For morphology and ALP stainings, the Axio Observer. A1 (Zeiss) was used.

##### *Widefield*

Immunofluorescence stainings were imaged with Axio Observer. Z1 (Zeiss) or the LAS X THUNDER (Leica). For live cell imaging of FURA-2-AM calcium imaging, an Olympus Motic AE31 microscope was used with an IonOptix epifluorescence acquisition system.

##### *Confocal*

For live cell imaging of Fluo-4-AM calcium imaging, the LSM710 confocal system (Zeiss) was used. For live cell imaging of TMRM/Mitotracker measurements the LAS X SP5 (Leica) was used.

##### *dSTORM*

7000 iPSC-CM were plated onto an eight-well chamber coverslip system (Cellvis). Cells were fixated 7 days later and stained with 1:500 RBM20 primary antibody and 1:500 Alexa-647 secondary antibody and a nucleus staining (DAPI). Instead of mounting solution, 500 µL 1x PBS was added to each well. The dSTORM imaging was performed using an Olympus IX-71 inverted microscope equipped with an APON 60xOTIRF (NA 1.49) oil immersion objective and a nosepiece stage (IX2-NPS, Olympus). Alexa Fluor 647 was excited using a 639 nm laser (Genesis MX639-1000, Coherent) at an intensity of approximately 2 $kW/cm^2$ to induce photoswitching. The imaging buffer consisted of PBS supplemented with 100 mM cysteamine hydrochloride (Sigma-Aldrich), pH 7.4, 4 % glucose, 8 units/ml glucose oxidase and 160 units/ml catalase. The excitation light was filtered through a 642/10 nm bandpass filter (Semrock) and focused on the back focal plane of the objective. A lens system and mirror, positioned on a linear translation stage, enabled switching between different illumination modes (EPI, HILO, and TIRF). Fluorescence emission was collected by the same objective and transmitted through a beam splitter (FF410/504/582/669-Di01, Semrock). The emitted light was further filtered using a bandpass filter (Brightline HC 679/41, Semrock) and projected onto an electron-multiplying CCD camera (iXon Ultra 897, Andor). Additional lenses in the detection path resulted in a final pixel size of 128.9 nm. A total of 15,000 frames were acquired at a frame rate of 67 Hz (15 ms exposure time). Image reconstruction was performed using rapi*d*STORM 3.3 [11], applying a threshold to exclude signals with fewer than 700 photons from the localization data. Further analysis was conducted using the software LOCAN [12]. To investigate RBM20 density within the nucleus, a region of interest (ROI) encompassing the nucleus was selected for each *d*STORM image. For analyzing RBM20 in the cytoplasm, an extended region around the nuclear ROI, with a distance of 2 µm, was chosen. For cluster analysis the localizations within these ROIs were grouped using a DBSCAN (Density-Based Spatial Clustering of Applications with Noise) [13] algorithm with parameters set to $\varepsilon = 20$ and minPoints = 3 [14]. A cluster was defined as a group of localizations with a minimum of three localizations, originating from one or a few secondary antibodies. Cluster densities in the nucleus and cytoplasm, as well as the ratio of these two densities (nucleus density : cytoplasm density) per cell were calculated.

#### Live cell imaging

##### *Calcium kinetics with Fluo-4-AM*

2.5 – 3 x $10^5$ iPSC-CM were plated onto Geltrex-coated 24 mm round coverslips and imaged 7-9 days later. Cells were loaded with 1.5 mL 1.8Tyrode solution (containing 1.8 mM calcium) containing 2,5 µM Fluo-4-AM and 0.025 % Pluronic (both from Thermo Fischer Scientific). Image acquisition with a Zeiss LSM 710 confocal microscope and a 63x 1.4 oil objective in line scan mode (512 pixels, 45.5 µm (zoom factor 3), 1057.7 Hz, 12 bit and 20,000 cycles at maximal speed). Calcium analysis was done as described by us previously for calcium kinetics and spark analysis with ImageJ Sparkmaster plugin [15]. For the spark analysis the 3D (width*height*amplitude) sum of a defined area during diastole was calculated. Drug treatments were conducted by directly diluting the compound into the measuring solution with the appropriate concentrations of Iso (1 µM), verapamil (30 nM) and metoprolol (5 µM). The drugs were incubated for 15 min before the subsequent measurement.

##### *Calcium content with FURA-2-AM*

3 – 4x$10^4$ iPSC-CMs were digested onto Geltrex – coated 35 mm round Fluoro dishes (World Precision Instruments) and imaged 7-9 days later. Cells were loaded with 5 µM FURA-2-AM dye was dissolved in 1 ml 1.25Tyrode solution (containing 1.25 mM calcium) for 15 min at 37 °C followed by incubation with 1.25Tyrode solution for 15 min at 37 °C. Finally, the 1.25Tyrode solution was refreshed and before measuring the cells. The FURA-2 loaded cells were excited at 340 and 380 nm and the emissions were measured at 510 nm. Data is shown as ratios of 340/380 nm with subtracted background values. For every Fluoro dish, 3-4 single cells were measured by recording $Ca^{2+}$ transients with increased pacing: 0.25, 0.5, 1.0, 2.0 and 3.0 Hz. The last measured cell was additionally stimulated with caffeine (10 µM) at the diastole. The parameters were analyzed using the IonOptix software.

##### *TMRM measurement*

2.5 – 3 x $10^5$ iPSC-CM were plated onto Geltrex-coated 24 mm round coverslips and imaged 7-9 days later. Cells were stained with 2.5 nM TMRM (Thermo Fisher Scientific) and 100 nM Mitotracker Green (Thermo Fisher Scientific) in Cardio Culture medium for 1 h at 37 °C. After 1 h , the solution was replaced with Cardio culture medium with 2.5 nM TMRM and images were acquired on the confocal system SP5 from Leica. For analysis, the mean value pixel intensity for the TMRM and Mitotracker channel were calculated using ImageJ and subsequently the ratio of TMRM/Mitotracker was used for quantification.

#### Seahorse

For simultaneous measurements of oxygen consumption rate (OCR), iPSC-CM were digested onto a Geltrex-coated 96-well cell plate (Seahorse, Agilent Billerica, MA, USA) at a density of 5 x $10^4$ cells per well. The cells were cultivated for 5-7 days under standard conditions as described above. Baseline respiration was measured in Seahorse XF base medium supplemented with 1 mM pyruvate and 5.5 mM glucose after calibration at 37 °C in an incubator without $CO_2$. Repeated measurements of OCR were performed over time. Metabolic states were measured after subsequent addition of 3 µM Oligomycin, 1 µM Carbonyl cyanide 4-(trifluoromethoxy)phenylhydrazone (FCCP), 3 µM Antimycin A, and 3 µM Rotenone. The data was normalized to protein amount using a BCA assay to correct for cell seeding differences.

#### Data analysis

##### *Sarcomere*

The sarcomeric regularity was analysis with FFT transformation, using the first peak height as a measurement of regularity [16].

##### *Mitochondrial Network*

Cells were incubated with 250 nM MitoSpy orange (Biolegend) for 30 min prior to fixation. After image acquisition, the mitochondrial network was analyzed using the ImageJ plugin MiNA as described previously [17].

##### *Statistics*

P-values are given for every significant result. The statistic test used is noted in each figure legend.


1. Rebs, S., et al., *Generation of pluripotent stem cell lines and CRISPR/Cas9 modified isogenic controls from a patient with dilated cardiomyopathy harboring a RBM20 p.R634W mutation.* Stem Cell Res, 2020. **47**: p. 101901.
2. Borchert, T., et al., *Catecholamine-Dependent beta-Adrenergic Signaling in a Pluripotent Stem Cell Model of Takotsubo Cardiomyopathy.* J Am Coll Cardiol, 2017. **70**(8): p. 975-991.
3. Tiburcy, M., et al., *Defined Engineered Human Myocardium With Advanced Maturation for Applications in Heart Failure Modeling and Repair.* Circulation, 2017. **135**(19): p. 1832-1847.
4. Tiburcy, M., et al., *Generation of Engineered Human Myocardium in a Multi-well Format.* STAR Protoc, 2020. **1**(1): p. 100032.
5. Chen, S., et al., *fastp: an ultra-fast all-in-one FASTQ preprocessor.* Bioinformatics, 2018. **34**(17): p. i884-i890.
6. Chen, S., *Ultrafast one-pass FASTQ data preprocessing, quality control, and deduplication using fastp.* Imeta, 2023. **2**(2): p. e107.
7. Liao, Y., G.K. Smyth, and W. Shi, *The Subread aligner: fast, accurate and scalable read mapping by seed-and-vote.* Nucleic Acids Res, 2013. **41**(10): p. e108.
8. Liao, Y., G.K. Smyth, and W. Shi, *featureCounts: an efficient general purpose program for assigning sequence reads to genomic features.* Bioinformatics, 2014. **30**(7): p. 923-30.
9. Love, M.I., W. Huber, and S. Anders, *Moderated estimation of fold change and dispersion for RNA-seq data with DESeq2.* Genome Biol, 2014. **15**(12): p. 550.
10. Anders, S., A. Reyes, and W. Huber, *Detecting differential usage of exons from RNA-seq data.* Genome Res, 2012. **22**(10): p. 2008-17.
11. Wolter, S., et al., *rapidSTORM: accurate, fast open-source software for localization microscopy.* Nat Methods, 2012. **9**(11): p. 1040-1.
12. Doose, S., *LOCAN: a python library for analyzing single-molecule localization microscopy data.* Bioinformatics, 2022. **38**(9): p. 2670-2672.
13. Schubert, E., et al., *DBSCAN Revisited, Revisited.* ACM Transactions on Database Systems, 2017. **42**(3): p. 1-21.
14. Ebert, V., et al., *Convex hull as diagnostic tool in single-molecule localization microscopy.* Bioinformatics, 2022. **38**(24): p. 5421-5429.
15. Haupt, L.P., et al., *Doxorubicin induces cardiotoxicity in a pluripotent stem cell model of aggressive B cell lymphoma cancer patients.* Basic Res Cardiol, 2022. **117**(1): p. 13.

16. Streckfuss-Bomeke, K., et al., *Severe DCM phenotype of patient harboring RBM20 mutation S635A can be modeled by patient-specific induced pluripotent stem cell-derived cardiomyocytes.* J Mol Cell Cardiol, 2017. **113**: p. 9-21.
17. Valente, A.J., et al., *A simple ImageJ macro tool for analyzing mitochondrial network morphology in mammalian cell culture.* Acta Histochem, 2017. **119**(3): p. 315-326.

#### Supplemental tables

***Suppl. Tab. 1: Summary of generated iPSC-lines from donor samples.***

| Patient donor | sample | Reprogramming | Intern-ID of clones |
|---|---|---|---|
| Healthy-1 from DCM II.3 | Skin/Fibroblasts | Sendai | S3RBM7<br>S3RBM8 |
| Healthy -2 from DCM III.3 | Skin/Fibroblasts | Sendai | S4RBM2<br>S4RBM6 |
| LVNC: II.3 | Skin/Fibroblasts | Plasmids | P6RBM1<br>P6RBM2 |
| DCM1: III.8 | Blood/PMBC | Sendai | S9RBM2<br>S9RBM4 |
| DCM2: III.10 | Skin/Fibroblasts | Sendai | S10RBM3<br>S10RBM5 |
| ResLVNC1 | Rescue in P6RBM1 | - | 6cr11<br>6cr20<br>6cr21<br>6cr29 |
| ResDCM1 | Rescue in S9RBM2 | - | 9cr35<br>9cr49 |
| LVNC-DCM.W | 634W mutation into 6cr29 | - | |

***Suppl. Tab. 2: List of antibodies used in this study.***

| Marker | Antibody | Dilution | ID and company |
|---|---|---|---|
| Pluripotency marker | IF: goat anti-OCT4 | 1:40 | R and D; AF1759<br>RRID AB_354975 |
| | IF: mouse anti-SOX2 | 1:200 | R and D; MAB2018<br>RRID AB_358009 |
| | IF: goat anti-LIN28 | 1:300 | R and D; AF3757<br>RRID AB_2234537 |
| | IF: goat anti-NANOG | 1:50 | R and D; AF1997<br>RRID AB_355097 |
| | IF: mouse anti-TRA1-60 | 1:200 | Abcam, ab16288<br>RRID AB_778563 |
| | IF: mouse anti-SSEA4 | 1:200 | Abcam, ab16287<br>RRID AB_778073 |

| | | | |
|---|---|---|---|
| Germlayer marker | IF: rabbit anti-AFP | 1:500 | Dako, A0008<br>RRID AB_2650473 |
| | IF: mouse anti-α-SMA | 1:3000 | Sigma, A2547<br>RRID AB_476701 |
| | IF: mouse anti-β-III-TUBULIN | 1:2000 | Covance, MMS-435P<br>RRID AB_2313773 |
| Sarcomeric marker | IF: mouse anti-α-actinin | 1:1000 | Sigma, A7811<br>RRID: AB_476766 |
| | IF: rabbit anti-Titin | 1:750 | MyoMedix, TTN-9<br>RRID:AB_2734750 |
| | FLOW: mouse anti-cTNT | 1:500 | Thermo Fisher, MS295PABX<br>RRID:AB_61810 |
| Cardiac function | IF: rabbit anti-RBM20 | 1:500 | Myomedix, RBM20-1 |
| | Western: mouse anti-PLN | 1:5000<br>in milk | Thermo Fisher, MA3-922<br>RRID:AB_2252716 |
| | Western: rabbit anti-PLN-S16p | 1:5000<br>in milk | Badrilla, A010-12AP<br>RRID:AB_2617047 |
| | Western: rabbit anti-PLN-T17p | 1:5000<br>in milk | Badrilla, A010-13<br>RRID:AB_2617048 |
| | Western: rabbit anti-RBM20 | 1:750<br>In BSA | Myomedix, RBM20-1 |
| Secondary antibody | IF: AF488 donkey anti-rabbit IgG | 1:1000 | Invitrogen, A31572<br>RRID AB_162543 |
| | IF: AF555 donkey anti-goat IgG | 1:1000 | Invitrogen, A21432<br>RRID AB_141788 |
| | IF: AF647 donkey anti-rabbit | 1:500 | Invitrogen, A31573<br>RRID:AB_2536183 |
| | IF and FLOW: AF488 donkey anti-mouse IgG | 1:1000 | Invitrogen, A21202<br>RRID AB_141607 |
| | IF: Cy3 goat anti-mouse IgG + IgM | 1:300 | Jackson Immuno, 115–165-068<br>RRID AB_2338686 |
| | Western:<br>ECL Mouse IgG HRP-Linked 1ml | 1:10000 | Th. Geyer, NA931<br>RRID:AB_772210 |
| | Western: ECL Rabbit IgG HRP-Linked 1ml | 1:10000 | Th. Geyer, NA934<br>RRID:AB_772206 |
| Nucleus staining | IF: Hoechst | 1:5000 | Sigma, 33258 |

***Suppl. Tab. 3: Primer list.***

| Gene | Forward 5´-> 3´ | Reverse 5´-> 3´ |
|---|---|---|
| 18s | ACCCGTTGAACCCCATTCGTGA | GCCTCACTAAACCATCCAATCGG |
| GAPDH | AGAGGCAGGGATGATGTTCT | TCTGCTGATGCCCCCATGTT |
| RBM20 – exon9 | GAGTGTACACAGTTACATGCAC | GTG GGACCTCGGGGAGA |
| RBM20 | CCTCCACTTGCCGCATATCTGT | AGACCAGGCATTTCTGAGCGTG |
| MYL2 | CGGAGAGGTTTTCCAAGGAGGA | CTCTTCTCCGTGGGTGATGATG |
| NR2F | CCGACCGGGTGGTCGCCTTTATGGA | CGGCTGGTTGGGGTACTGGCTCCTA |
| TTN-N2B | CCAATGAGTATGGCAGTGTCA | TACGTTCCGGAAGTAATTTGC |
| TTN-N2BA | GCCACACTAACTGTGACAGAGG | GGCTGCCTTACCCACAAAAG |
| RYR2-24bp | GTCACAGGATCCCAACGCAG | CTTTGCTGGCACTGATTGTCTG |
| RYR2 | CTTGAGGTTGGCTTTCTGCCAG | TGTGCCAGCAAAGAGAGGAGAC |
| LDB3-exon5 | TCAAAGCGTCCCATTCCCATC | CGGGAGAA GCAGGGCTAAA |
| LDB3 | ACCTCGTGGTGGCCATTG | GTGGAGATGGGAATGGGACG |
| CAMk2D-exon14 | CCATCTTGACAACTATGCTGGCT | GAACACTCGAACTGGACTTCCT |
| CAMk2D | ACACGGTGACTCCTGAAGCCAA | GTCTCCTGTCTGTGCATCATGG |
| TRDN-exon9 | GTCCATGGGGATTTAAAACCAGG | CTTCAAGGGCAGGTGATGC |
| TRDN | GGAGGACAAAGAGAAAGCAGCTG | AGGTGGAATGGCTGGGCTTTGT |
| IMMT-exon5-6 | AAACAGCCTGCCTCACAACT | TCCTTCAATGCACCCTCCAC |
| IMMT | CAGGCTGTCAATGCACACTCCA | CATCTACTGCCTTTCTGCGTTCC |

***Suppl. Tab. 4: Top 100 differentially expressed genes resLVNC vs LVNC.***

| ENTREZID | SYMBOL | GENENAME | Log2 Fold Change | Raw P-value | Adjusted P-value |
|---|---|---|---|---|---|
| 358 | AQP1 | aquaporin 1 (Colton blood group) | -1.951±0.238 | 6.32 × 10^-16 | 1.30 × 10^-11 |
| 3959 | LGALS3BP | galectin 3 binding protein | -2.607±0.338 | 4.99 × 10^-14 | 3.52 × 10^-10 |
| 102724652 | LOC102724652 | crystallin alpha A2 | -0.756±0.804 | 6.79 × 10^-14 | 3.52 × 10^-10 |
| 1409 | CRYAA | crystallin alpha A | -0.754±0.804 | 6.83 × 10^-14 | 3.52 × 10^-10 |
| 112 | ADCY6 | adenylate cyclase 6 | -1.882±0.256 | 4.54 × 10^-13 | 1.87 × 10^-9 |
| 26001 | RNF167 | ring finger protein 167 | -1.402±0.193 | 6.01 × 10^-13 | 2.07 × 10^-9 |
| 80023 | NRSN2 | neurensin 2 | -2.705±0.373 | 1.09 × 10^-12 | 3.20 × 10^-9 |
| 3949 | LDLR | low density lipoprotein receptor | -2.188±0.308 | 2.37 × 10^-12 | 6.10 × 10^-9 |
| 37 | ACADVL | acyl-CoA dehydrogenase very long chain | -1.572±0.224 | 4.33 × 10^-12 | 9.93 × 10^-9 |
| 6840 | SVIL | supervillin | -0.945±0.137 | 6.11 × 10^-12 | 1.15 × 10^-8 |
| 8048 | CSRP3 | cysteine and glycine rich protein 3 | -1.743±0.250 | 6.04 × 10^-12 | 1.15 × 10^-8 |
| 6604 | SMARCD3 | SWI/SNF related, matrix associated, actin dependent regulator of chromatin, subfamily d, member 3 | -1.552±0.223 | 6.73 × 10^-12 | 1.16 × 10^-8 |

| ENTREZID | SYMBOL | GENENAME | Log2 Fold Change | Raw P-value | Adjusted P-value |
|---|---|---|---|---|---|
| 11180 | WDR6 | WD repeat domain 6 | -2.405±0.342 | 7.39 × 10^-12 | 1.17 × 10^-8 |
| 327 | APEH | acylaminoacyl-peptide hydrolase | -2.713±0.395 | 2.12 × 10^-11 | 3.12 × 10^-8 |
| 2044 | EPHA5 | EPH receptor A5 | 4.556±0.641 | 2.87 × 10^-11 | 3.94 × 10^-8 |
| 7284 | TUFM | Tu translation elongation factor, mitochondrial | -2.311±0.357 | 1.81 × 10^-10 | 2.12 × 10^-7 |
| 6711 | SPTBN1 | spectrin beta, non-erythrocytic 1 | -1.223±0.189 | 1.85 × 10^-10 | 2.12 × 10^-7 |
| 10221 | TRIB1 | tribbles pseudokinase 1 | -1.661±0.257 | 1.71 × 10^-10 | 2.12 × 10^-7 |
| 1523 | CUX1 | cut like homeobox 1 | -1.108±0.173 | 2.05 × 10^-10 | 2.22 × 10^-7 |
| 4240 | MFGE8 | milk fat globule EGF and factor V/VIII domain containing | -2.654±0.407 | 2.56 × 10^-10 | 2.64 × 10^-7 |
| 7057 | THBS1 | thrombospondin 1 | -1.927±0.302 | 3.81 × 10^-10 | 3.74 × 10^-7 |
| 286336 | FAM78A | family with sequence similarity 78 member A | -1.999±0.314 | 4.17 × 10^-10 | 3.91 × 10^-7 |
| 25805 | BAMBI | BMP and activin membrane bound inhibitor | -1.781±0.282 | 5.41 × 10^-10 | 4.85 × 10^-7 |
| 163702 | IFNLR1 | interferon lambda receptor 1 | -2.356±0.375 | 6.12 × 10^-10 | 5.26 × 10^-7 |
| 257194 | NEGR1 | neuronal growth regulator 1 | 2.033±0.326 | 7.14 × 10^-10 | 5.89 × 10^-7 |
| 3693 | ITGB5 | integrin subunit beta 5 | -1.973±0.318 | 9.20 × 10^-10 | 7.30 × 10^-7 |
| 30845 | EHD3 | EH domain containing 3 | -1.924±0.310 | 1.16 × 10^-9 | 8.86 × 10^-7 |
| 196385 | DNAH10 | dynein axonemal heavy chain 10 | 2.718±0.433 | 1.60 × 10^-9 | 1.14 × 10^-6 |
| 593 | BCKDHA | branched chain keto acid dehydrogenase E1 subunit alpha | -2.845±0.453 | 1.60 × 10^-9 | 1.14 × 10^-6 |
| 826 | CAPNS1 | calpain small subunit 1 | -1.693±0.275 | 1.66 × 10^-9 | 1.14 × 10^-6 |
| 146691 | TOM1L2 | target of myb1 like 2 membrane trafficking protein | -1.667±0.273 | 1.79 × 10^-9 | 1.15 × 10^-6 |
| 58498 | MYL7 | myosin light chain 7 | -2.473±0.395 | 1.77 × 10^-9 | 1.15 × 10^-6 |
| 27122 | DKK3 | dickkopf WNT signaling pathway inhibitor 3 | -1.347±0.223 | 2.83 × 10^-9 | 1.77 × 10^-6 |
| 6506 | SLC1A2 | solute carrier family 1 member 2 | 2.574±0.421 | 3.13 × 10^-9 | 1.85 × 10^-6 |
| 2896 | GRN | granulin precursor | -3.089±0.500 | 3.06 × 10^-9 | 1.85 × 10^-6 |
| 3679 | ITGA7 | integrin subunit alpha 7 | -2.036±0.338 | 3.27 × 10^-9 | 1.85 × 10^-6 |

| ENTREZID | SYMBOL | GENENAME | Log2 Fold Change | Raw P-value | Adjusted P-value |
|---|---|---|---|---|---|
| 65997 | RASL11B | RAS like family 11 member B | -1.859±0.310 | 3.32 × 10^-9 | 1.85 × 10^-6 |
| 55089 | SLC38A4 | solute carrier family 38 member 4 | 2.996±0.496 | 3.42 × 10^-9 | 1.86 × 10^-6 |
| 1917 | EEF1A2 | eukaryotic translation elongation factor 1 alpha 2 | -2.815±0.462 | 3.92 × 10^-9 | 2.07 × 10^-6 |
| 200162 | SPAG17 | sperm associated antigen 17 | 4.466±0.686 | 4.44 × 10^-9 | 2.26 × 10^-6 |
| 57658 | CALCOCO1 | calcium binding and coiled-coil domain 1 | -1.204±0.203 | 4.48 × 10^-9 | 2.26 × 10^-6 |
| 4607 | MYBPC3 | myosin binding protein C3 | -2.193±0.362 | 4.73 × 10^-9 | 2.32 × 10^-6 |
| 7436 | VLDLR | very low density lipoprotein receptor | -1.059±0.180 | 5.31 × 10^-9 | 2.55 × 10^-6 |
| 4192 | MDK | midkine | -1.866±0.311 | 5.51 × 10^-9 | 2.55 × 10^-6 |
| 57158 | JPH2 | junctophilin 2 | -1.832±0.308 | 5.56 × 10^-9 | 2.55 × 10^-6 |
| 3213 | HOXB3 | homeobox B3 | 1.997±0.340 | 8.83 × 10^-9 | 3.96 × 10^-6 |
| 1284 | COL4A2 | collagen type IV alpha 2 chain | -2.368±0.402 | 1.25 × 10^-8 | 5.47 × 10^-6 |
| 65018 | PINK1 | PTEN induced kinase 1 | -2.786±0.467 | 1.32 × 10^-8 | 5.67 × 10^-6 |
| 146862 | UNC45B | unc-45 myosin chaperone B | -1.374±0.238 | 1.37 × 10^-8 | 5.75 × 10^-6 |
| 2975 | GTF3C1 | general transcription factor IIIC subunit 1 | -1.249±0.219 | 1.67 × 10^-8 | 6.90 × 10^-6 |
| 94005 | PIGS | phosphatidylinositol glycan anchor biosynthesis class S | -1.589±0.281 | 2.15 × 10^-8 | 8.40 × 10^-6 |
| 4924 | NUCB1 | nucleobindin 1 | -2.381±0.411 | 2.16 × 10^-8 | 8.40 × 10^-6 |
| 7074 | TIAM1 | TIAM Rac1 associated GEF 1 | 2.195±0.382 | 2.10 × 10^-8 | 8.40 × 10^-6 |
| 30008 | EFEMP2 | EGF containing fibulin extracellular matrix protein 2 | -2.143±0.377 | 2.26 × 10^-8 | 8.65 × 10^-6 |
| 4258 | MGST2 | microsomal glutathione S-transferase 2 | -1.423±0.252 | 2.41 × 10^-8 | 9.05 × 10^-6 |
| 9844 | ELMO1 | engulfment and cell motility 1 | 2.221±0.387 | 2.66 × 10^-8 | 9.78 × 10^-6 |
| 777 | CACNA1E | calcium voltage-gated channel subunit alpha1 E | 3.955±0.659 | 2.86 × 10^-8 | 1.04 × 10^-5 |
| 147906 | DACT3 | dishevelled binding antagonist of beta catenin 3 | -1.751±0.313 | 3.14 × 10^-8 | 1.12 × 10^-5 |
| 23467 | NPTXR | neuronal pentraxin receptor | -1.689±0.300 | 3.25 × 10^-8 | 1.14 × 10^-5 |
| 8828 | NRP2 | neuropilin 2 | -1.377±0.247 | 3.37 × 10^-8 | 1.16 × 10^-5 |

| ENTREZID | SYMBOL | GENENAME | Log2 Fold Change | Raw P-value | Adjusted P-value |
|---|---|---|---|---|---|
| 9348 | NDST3 | N-deacetylase and N-sulfotransferase 3 | 3.872±0.646 | 3.45 × 10^-8 | 1.17 × 10^-5 |
| 3913 | LAMB2 | laminin subunit beta 2 | -2.071±0.363 | 3.66 × 10^-8 | 1.22 × 10^-5 |
| 115123 | MARCHF3 | membrane associated ring-CH-type finger 3 | -1.640±0.298 | 4.21 × 10^-8 | 1.38 × 10^-5 |
| 79915 | ATAD5 | ATPase family AAA domain containing 5 | 1.722±0.310 | 4.37 × 10^-8 | 1.41 × 10^-5 |
| 3673 | ITGA2 | integrin subunit alpha 2 | -1.838±0.338 | 4.48 × 10^-8 | 1.42 × 10^-5 |
| 8566 | PDXK | pyridoxal kinase | -1.529±0.276 | 4.65 × 10^-8 | 1.45 × 10^-5 |
| 5156 | PDGFRA | platelet derived growth factor receptor alpha | -1.898±0.345 | 4.81 × 10^-8 | 1.48 × 10^-5 |
| 5922 | RASA2 | RAS p21 protein activator 2 | 1.515±0.277 | 5.15 × 10^-8 | 1.56 × 10^-5 |
| 23038 | WDTC1 | WD and tetratricopeptide repeats 1 | -2.258±0.412 | 5.70 × 10^-8 | 1.70 × 10^-5 |
| 10723 | SLC12A7 | solute carrier family 12 member 7 | -3.345±0.569 | 6.20 × 10^-8 | 1.83 × 10^-5 |
| 51555 | PEX5L | peroxisomal biogenesis factor 5 like | 2.645±0.470 | 6.50 × 10^-8 | 1.89 × 10^-5 |
| 10622 | POLR3G | RNA polymerase III subunit G | 2.255±0.408 | 7.05 × 10^-8 | 2.02 × 10^-5 |
| 2621 | GAS6 | growth arrest specific 6 | -2.942±0.521 | 7.15 × 10^-8 | 2.02 × 10^-5 |
| 8626 | TP63 | tumor protein p63 | 3.631±0.619 | 7.61 × 10^-8 | 2.12 × 10^-5 |
| 219595 | FOLH1B | folate hydrolase 1B (pseudogene) | 3.548±0.623 | 7.96 × 10^-8 | 2.19 × 10^-5 |
| 2846 | LPAR4 | lysophosphatidic acid receptor 4 | 3.554±0.617 | 8.07 × 10^-8 | 2.19 × 10^-5 |
| 6900 | CNTN2 | contactin 2 | 4.292±0.734 | 8.81 × 10^-8 | 2.36 × 10^-5 |
| 672 | BRCA1 | BRCA1 DNA repair associated | 1.768±0.328 | 8.93 × 10^-8 | 2.36 × 10^-5 |
| 5339 | PLEC | plectin | -2.342±0.423 | 9.48 × 10^-8 | 2.48 × 10^-5 |
| 9469 | CHST3 | carbohydrate sulfotransferase 3 | -1.829±0.338 | 9.69 × 10^-8 | 2.50 × 10^-5 |
| 10690 | FUT9 | fucosyltransferase 9 | 3.495±0.619 | 1.03 × 10^-7 | 2.61 × 10^-5 |
| 29106 | SCG3 | secretogranin III | 3.862±0.668 | 1.09 × 10^-7 | 2.74 × 10^-5 |
| 25844 | YIPF3 | Yip1 domain family member 3 | -1.948±0.361 | 1.12 × 10^-7 | 2.79 × 10^-5 |
| 5819 | NECTIN2 | nectin cell adhesion molecule 2 | -2.739±0.501 | 1.17 × 10^-7 | 2.86 × 10^-5 |

| ENTREZID | SYMBOL | GENENAME | Log2 Fold Change | Raw P-value | Adjusted P-value |
|---|---|---|---|---|---|
| 389 | RHOC | ras homolog family member C | -2.059±0.378 | 1.20 × 10^-7 | 2.89 × 10^-5 |
| 5315 | PKM | pyruvate kinase M1/2 | -1.854±0.340 | 1.19 × 10^-7 | 2.89 × 10^-5 |
| 2137 | EXTL3 | exostosin like glycosyltransferase 3 | -1.296±0.243 | 1.33 × 10^-7 | 3.16 × 10^-5 |
| 7168 | TPM1 | tropomyosin 1 | -1.529±0.283 | 1.49 × 10^-7 | 3.49 × 10^-5 |
| 105375661 | NA | NA | 5.129±0.864 | 1.54 × 10^-7 | 3.58 × 10^-5 |
| 23385 | NCSTN | nicastrin | -1.005±0.190 | 1.60 × 10^-7 | 3.66 × 10^-5 |
| 84457 | PHYHIPL | phytanoyl-CoA 2-hydroxylase interacting protein like | 3.390±0.599 | 1.63 × 10^-7 | 3.69 × 10^-5 |
| 58476 | TP53INP2 | tumor protein p53 inducible nuclear protein 2 | -1.388±0.262 | 1.65 × 10^-7 | 3.71 × 10^-5 |
| 5454 | POU3F2 | POU class 3 homeobox 2 | 5.074±0.839 | 1.69 × 10^-7 | 3.75 × 10^-5 |
| 91304 | TMEM259 | transmembrane protein 259 | -2.016±0.375 | 1.73 × 10^-7 | 3.79 × 10^-5 |
| 56981 | PRDM11 | PR/SET domain 11 | -1.270±0.242 | 1.78 × 10^-7 | 3.87 × 10^-5 |
| 339123 | JMJD8 | jumonji domain containing 8 | -3.411±0.598 | 1.88 × 10^-7 | 4.03 × 10^-5 |
| 558 | AXL | AXL receptor tyrosine kinase | -1.991±0.379 | 1.91 × 10^-7 | 4.07 × 10^-5 |
| 7005 | TEAD3 | TEA domain transcription factor 3 | -2.006±0.377 | 2.07 × 10^-7 | 4.35 × 10^-5 |
| 3636 | INPPL1 | inositol polyphosphate phosphatase like 1 | -1.666±0.313 | 2.40 × 10^-7 | 5.00 × 10^-5 |
| 780 | DDR1 | discoidin domain receptor tyrosine kinase 1 | -2.286±0.426 | 2.52 × 10^-7 | 5.09 × 10^-5 |

***Suppl. Tab. 4: Top 100 differentially expressed genes resDCM1 vs DCM1.***

| ENTREZID | SYMBOL | GENENAME | Log2 Fold Change | Raw P-value | Adjusted P-value |
|---|---|---|---|---|---|
| 10399 | RACK1 | receptor for activated C kinase 1 | -2.185±0.116 | 4.01 × 10^-79 | 7.99 × 10^-75 |
| 6134 | RPL10 | ribosomal protein L10 | -2.533±0.139 | 4.09 × 10^-74 | 4.07 × 10^-70 |
| 1937 | EEF1G | eukaryotic translation elongation factor 1 gamma | -2.088±0.119 | 1.82 × 10^-68 | 1.21 × 10^-64 |
| 7316 | UBC | ubiquitin C | -2.474±0.148 | 2.34 × 10^-62 | 1.16 × 10^-58 |
| 230 | ALDOC | aldolase, fructose-bisphosphate C | -3.683±0.237 | 1.21 × 10^-53 | 4.81 × 10^-50 |

| ENTREZID | SYMBOL | GENENAME | Log2 Fold Change | Raw P-value | Adjusted P-value |
|---|---|---|---|---|---|
| 728658 | RPL13AP5 | ribosomal protein L13a pseudogene 5 | -2.172±0.146 | 9.01 × 10^-50 | 2.99 × 10^-46 |
| 967 | CD63 | CD63 molecule | -2.905±0.209 | 5.61 × 10^-43 | 1.60 × 10^-39 |
| 6202 | RPS8 | ribosomal protein S8 | -1.717±0.125 | 2.06 × 10^-42 | 5.13 × 10^-39 |
| 6143 | RPL19 | ribosomal protein L19 | -1.668±0.129 | 2.88 × 10^-38 | 6.38 × 10^-35 |
| 4637 | MYL6 | myosin light chain 6 | -1.778±0.141 | 4.23 × 10^-36 | 8.43 × 10^-33 |
| 6208 | RPS14 | ribosomal protein S14 | -2.504±0.197 | 5.80 × 10^-36 | 1.05 × 10^-32 |
| 498 | ATP5F1A | ATP synthase F1 subunit alpha | -1.588±0.128 | 1.62 × 10^-35 | 2.68 × 10^-32 |
| 3959 | LGALS3BP | galectin 3 binding protein | -3.136±0.251 | 5.36 × 10^-35 | 8.21 × 10^-32 |
| 1642 | DDB1 | damage specific DNA binding protein 1 | -1.419±0.115 | 1.11 × 10^-34 | 1.58 × 10^-31 |
| 5213 | PFKM | phosphofructokinase, muscle | -2.462±0.200 | 1.52 × 10^-34 | 2.01 × 10^-31 |
| 10939 | AFG3L2 | AFG3 like matrix AAA peptidase subunit 2 | -1.598±0.132 | 1.05 × 10^-33 | 1.31 × 10^-30 |
| 800 | CALD1 | caldesmon 1 | 1.738±0.145 | 3.72 × 10^-33 | 4.35 × 10^-30 |
| 9669 | EIF5B | eukaryotic translation initiation factor 5B | 2.253±0.188 | 4.53 × 10^-33 | 5.01 × 10^-30 |
| 4720 | NDUFS2 | NADH:ubiquinone oxidoreductase core subunit S2 | -1.894±0.159 | 1.54 × 10^-32 | 1.62 × 10^-29 |
| 3945 | LDHB | lactate dehydrogenase B | -1.929±0.164 | 6.91 × 10^-32 | 6.88 × 10^-29 |
| 6130 | RPL7A | ribosomal protein L7a | -2.125±0.181 | 2.28 × 10^-31 | 2.16 × 10^-28 |
| 4541 | ND6 | NADH dehydrogenase subunit 6 | 3.950±0.337 | 2.64 × 10^-31 | 2.39 × 10^-28 |
| 5250 | SLC25A3 | solute carrier family 25 member 3 | -1.654±0.145 | 4.05 × 10^-30 | 3.51 × 10^-27 |
| 7415 | VCP | valosin containing protein | -1.926±0.171 | 2.06 × 10^-29 | 1.71 × 10^-26 |
| 6132 | RPL8 | ribosomal protein L8 | -2.745±0.243 | 6.39 × 10^-29 | 5.09 × 10^-26 |
| 6415 | SELENOW | selenoprotein W | -3.895±0.343 | 8.37 × 10^-29 | 6.41 × 10^-26 |
| 6122 | RPL3 | ribosomal protein L3 | -1.926±0.173 | 1.10 × 10^-28 | 8.10 × 10^-26 |
| 374393 | FAM111B | FAM111 trypsin like peptidase B | 2.504±0.226 | 1.42 × 10^-28 | 1.01 × 10^-25 |
| 388524 | RPSA2 | ribosomal protein SA 2 | -2.141±0.194 | 4.01 × 10^-28 | 2.75 × 10^-25 |

| ENTREZID | SYMBOL | GENENAME | Log2 Fold Change | Raw P-value | Adjusted P-value |
|---|---|---|---|---|---|
| 63967 | CLSPN | claspin | 1.961±0.179 | 7.10 × 10^-28 | 4.56 × 10^-25 |
| 4678 | NASP | nuclear autoantigenic sperm protein | 1.249±0.114 | 7.03 × 10^-28 | 4.56 × 10^-25 |
| 9500 | MAGED1 | MAGE family member D1 | -2.188±0.201 | 1.90 × 10^-27 | 1.18 × 10^-24 |
| 1499 | CTNNB1 | catenin beta 1 | -1.549±0.143 | 3.73 × 10^-27 | 2.25 × 10^-24 |
| 6175 | RPLP0 | ribosomal protein lateral stalk subunit P0 | -1.525±0.142 | 1.17 × 10^-26 | 6.83 × 10^-24 |
| 2170 | FABP3 | fatty acid binding protein 3 | -1.571±0.148 | 2.26 × 10^-26 | 1.29 × 10^-23 |
| 653513 | LOC653513 | phosphodiesterase 4D interacting protein-like | -2.169±0.204 | 3.28 × 10^-26 | 1.82 × 10^-23 |
| 23326 | USP22 | ubiquitin specific peptidase 22 | -1.220±0.117 | 2.20 × 10^-25 | 1.19 × 10^-22 |
| 3921 | RPSA | ribosomal protein SA | -2.601±0.247 | 3.17 × 10^-25 | 1.66 × 10^-22 |
| 1340 | COX6B1 | cytochrome c oxidase subunit 6B1 | -1.512±0.146 | 3.53 × 10^-25 | 1.80 × 10^-22 |
| 506 | ATP5F1B | ATP synthase F1 subunit beta | -1.563±0.152 | 1.11 × 10^-24 | 5.55 × 10^-22 |
| 84081 | NSRP1 | nuclear speckle splicing regulatory protein 1 | 1.765±0.172 | 1.27 × 10^-24 | 6.15 × 10^-22 |
| 1345 | COX6C | cytochrome c oxidase subunit 6C | -1.428±0.139 | 1.60 × 10^-24 | 7.60 × 10^-22 |
| 5660 | PSAP | prosaposin | -2.357±0.230 | 1.99 × 10^-24 | 9.23 × 10^-22 |
| 25824 | PRDX5 | peroxiredoxin 5 | -3.232±0.308 | 3.36 × 10^-24 | 1.52 × 10^-21 |
| 118 | ADD1 | adducin 1 | -1.488±0.147 | 4.45 × 10^-24 | 1.97 × 10^-21 |
| 10574 | CCT7 | chaperonin containing TCP1 subunit 7 | -1.571±0.155 | 6.99 × 10^-24 | 3.03 × 10^-21 |
| 517 | ATP5MC2 | ATP synthase membrane subunit c locus 2 | -2.211±0.219 | 9.40 × 10^-24 | 3.99 × 10^-21 |
| 1666 | DECR1 | 2,4-dienoyl-CoA reductase 1 | -1.791±0.178 | 1.06 × 10^-23 | 4.39 × 10^-21 |
| 5858 | PZP | PZP alpha-2-macroglobulin like | 2.820±0.281 | 1.71 × 10^-23 | 6.93 × 10^-21 |
| 23107 | MRPS27 | mitochondrial ribosomal protein S27 | -2.372±0.236 | 2.62 × 10^-23 | 1.05 × 10^-20 |
| 3032 | HADHB | hydroxyacyl-CoA dehydrogenase trifunctional multienzyme complex subunit beta | -1.385±0.140 | 6.07 × 10^-23 | 2.37 × 10^-20 |
| 309 | ANXA6 | annexin A6 | -1.379±0.140 | 8.71 × 10^-23 | 3.34 × 10^-20 |
| 3006 | H1-2 | H1.2 linker histone, cluster member | -2.927±0.292 | 8.89 × 10^-23 | 3.34 × 10^-20 |

| ENTREZID | SYMBOL | GENENAME | Log2 Fold Change | Raw P-value | Adjusted P-value |
|---|---|---|---|---|---|
| 4538 | ND4 | NADH dehydrogenase subunit 4 | 2.711±0.275 | 9.77 × 10^-23 | 3.61 × 10^-20 |
| 10541 | ANP32B | acidic nuclear phosphoprotein 32 family member B | 2.222±0.227 | 1.09 × 10^-22 | 3.95 × 10^-20 |
| 5430 | POLR2A | RNA polymerase II subunit A | 1.908±0.195 | 1.12 × 10^-22 | 3.99 × 10^-20 |
| 23521 | RPL13A | ribosomal protein L13a | -1.510±0.153 | 1.20 × 10^-22 | 4.19 × 10^-20 |
| 150082 | LCA5L | lebercilin LCA5 like | 3.067±0.311 | 1.30 × 10^-22 | 4.46 × 10^-20 |
| 284459 | ZNF875 | zinc finger protein 875 | -2.032±0.206 | 1.43 × 10^-22 | 4.82 × 10^-20 |
| 10106 | CTDSP2 | CTD small phosphatase 2 | -1.443±0.148 | 1.80 × 10^-22 | 5.96 × 10^-20 |
| 195828 | ZNF367 | zinc finger protein 367 | 1.524±0.157 | 2.30 × 10^-22 | 7.51 × 10^-20 |
| 37 | ACADVL | acyl-CoA dehydrogenase very long chain | -1.642±0.169 | 2.55 × 10^-22 | 8.21 × 10^-20 |
| 6136 | RPL12 | ribosomal protein L12 | -1.723±0.177 | 2.75 × 10^-22 | 8.68 × 10^-20 |
| 29896 | TRA2A | transformer 2 alpha homolog | 1.381±0.143 | 3.66 × 10^-22 | 1.14 × 10^-19 |
| 2137 | EXTL3 | exostosin like glycosyltransferase 3 | -2.093±0.216 | 4.63 × 10^-22 | 1.42 × 10^-19 |
| 8367 | H4C5 | H4 clustered histone 5 | -2.407±0.246 | 4.89 × 10^-22 | 1.48 × 10^-19 |
| 389247 | ING2-DT | ING2 divergent transcript | 4.380±0.451 | 1.07 × 10^-21 | 3.17 × 10^-19 |
| 2030 | SLC29A1 | solute carrier family 29 member 1 (Augustine blood group) | -2.442±0.253 | 1.28 × 10^-21 | 3.75 × 10^-19 |
| 4190 | MDH1 | malate dehydrogenase 1 | -1.312±0.137 | 1.44 × 10^-21 | 4.16 × 10^-19 |
| 6774 | STAT3 | signal transducer and activator of transcription 3 | -1.334±0.140 | 1.71 × 10^-21 | 4.87 × 10^-19 |
| 7314 | UBB | ubiquitin B | -1.216±0.128 | 1.87 × 10^-21 | 5.24 × 10^-19 |
| 5702 | PSMC3 | proteasome 26S subunit, ATPase 3 | -2.243±0.234 | 2.05 × 10^-21 | 5.68 × 10^-19 |
| 54892 | NCAPG2 | non-SMC condensin II complex subunit G2 | 1.680±0.176 | 2.13 × 10^-21 | 5.81 × 10^-19 |
| 10476 | ATP5PD | ATP synthase peripheral stalk subunit d | -2.236±0.235 | 4.50 × 10^-21 | 1.21 × 10^-18 |
| 55215 | FANCI | FA complementation group I | 1.595±0.169 | 5.33 × 10^-21 | 1.42 × 10^-18 |
| 1072 | CFL1 | cofilin 1 | -1.465±0.156 | 9.57 × 10^-21 | 2.51 × 10^-18 |
| 3073 | HEXA | hexosaminidase subunit alpha | -2.262±0.238 | 1.60 × 10^-20 | 4.15 × 10^-18 |

| ENTREZID | SYMBOL | GENENAME | Log2 Fold Change | Raw P-value | Adjusted P-value |
|---|---|---|---|---|---|
| 5518 | PPP2R1A | protein phosphatase 2 scaffold subunit Aalpha | -2.462±0.265 | 3.05 × 10^-20 | 7.78 × 10^-18 |
| 284361 | EMC10 | ER membrane protein complex subunit 10 | 2.196±0.239 | 3.19 × 10^-20 | 8.04 × 10^-18 |
| 161497 | STRC | stereocilin | 3.670±0.394 | 4.00 × 10^-20 | 9.83 × 10^-18 |
| 1327 | COX4I1 | cytochrome c oxidase subunit 4I1 | -2.252±0.244 | 3.95 × 10^-20 | 9.83 × 10^-18 |
| 105378510 | NA | NA | 3.944±0.427 | 5.93 × 10^-20 | 1.42 × 10^-17 |
| 9168 | TMSB10 | thymosin beta 10 | -1.659±0.181 | 5.97 × 10^-20 | 1.42 × 10^-17 |
| 27000 | DNAJC2 | DnaJ heat shock protein family (Hsp40) member C2 | 1.434±0.157 | 5.90 × 10^-20 | 1.42 × 10^-17 |
| 4509 | ATP8 | ATP synthase F0 subunit 8 | 2.997±0.329 | 1.22 × 10^-19 | 2.85 × 10^-17 |
| 8031 | NCOA4 | nuclear receptor coactivator 4 | -1.605±0.177 | 1.59 × 10^-19 | 3.68 × 10^-17 |
| 4539 | ND4L | NADH dehydrogenase subunit 4L | 2.440±0.269 | 1.67 × 10^-19 | 3.83 × 10^-17 |
| 6567 | SLC16A2 | solute carrier family 16 member 2 | -2.574±0.283 | 2.21 × 10^-19 | 5.00 × 10^-17 |
| 10916 | MAGED2 | MAGE family member D2 | -1.939±0.215 | 2.78 × 10^-19 | 6.22 × 10^-17 |
| 220869 | ZNG1E | Zn regulated GTPase metalloprotein activator 1E | 1.305±0.146 | 3.15 × 10^-19 | 6.97 × 10^-17 |
| 6155 | RPL27 | ribosomal protein L27 | -1.251±0.140 | 4.26 × 10^-19 | 9.33 × 10^-17 |
| 4540 | ND5 | NADH dehydrogenase subunit 5 | 2.769±0.308 | 4.59 × 10^-19 | 9.94 × 10^-17 |
| 2934 | GSN | gelsolin | -1.993±0.223 | 5.23 × 10^-19 | 1.12 × 10^-16 |
| 343099 | CCDC18 | coiled-coil domain containing 18 | 1.693±0.190 | 6.03 × 10^-19 | 1.28 × 10^-16 |
| 6141 | RPL18 | ribosomal protein L18 | -2.350±0.260 | 6.74 × 10^-19 | 1.41 × 10^-16 |
| 8508 | NIPSNAP1 | nipsnap homolog 1 | -2.286±0.254 | 6.85 × 10^-19 | 1.42 × 10^-16 |
| 6185 | RPN2 | ribophorin II | -1.588±0.178 | 6.96 × 10^-19 | 1.43 × 10^-16 |
| 284695 | ZNF326 | zinc finger protein 326 | 1.614±0.182 | 8.01 × 10^-19 | 1.63 × 10^-16 |
| 1938 | EEF2 | eukaryotic translation elongation factor 2 | -0.928±0.105 | 9.17 × 10^-19 | 1.85 × 10^-16 |
| 8899 | PRPF4B | pre-mRNA processing factor 4B | 1.401±0.159 | 9.67 × 10^-19 | 1.93 × 10^-16 |

 ***Suppl. Tab. 6: Top 24 differentially expressed genes LVNC vs DCM1.***

| ENTREZID | SYMBOL | GENENAME | Log2 Fold Change | Raw P-value | Adjusted P-value |
|---|---|---|---|---|---|
| 8287 | USP9Y | ubiquitin specific peptidase 9 Y-linked | -7.291±0.597 | 1.28 × 10^-21 | 2.44 × 10^-17 |
| 8653 | DDX3Y | DEAD-box helicase 3 Y-linked | -7.411±0.607 | 2.42 × 10^-20 | 2.30 × 10^-16 |
| 7404 | UTY | ubiquitously transcribed tetratricopeptide repeat containing, Y-linked | -6.797±0.650 | 1.94 × 10^-16 | 1.23 × 10^-12 |
| 6192 | RPS4Y1 | ribosomal protein S4 Y-linked 1 | -6.183±0.666 | 2.64 × 10^-14 | 1.26 × 10^-10 |
| 8284 | KDM5D | lysine demethylase 5D | -6.096±0.671 | 4.93 × 10^-14 | 1.87 × 10^-10 |
| 9086 | EIF1AY | eukaryotic translation initiation factor 1A Y-linked | -6.069±0.676 | 7.59 × 10^-14 | 2.41 × 10^-10 |
| 7544 | ZFY | zinc finger protein Y-linked | -5.719±0.685 | 8.24 × 10^-13 | 2.24 × 10^-9 |
| 84171 | LOXL4 | lysyl oxidase like 4 | -3.285±0.494 | 3.87 × 10^-11 | 9.21 × 10^-8 |
| 246126 | TXLNGY | taxilin gamma Y-linked (pseudogene) | -5.205±0.704 | 5.54 × 10^-11 | 1.17 × 10^-7 |
| 64595 | TTTY15 | testis expressed transcript, Y-linked 15 | -4.786±0.723 | 8.51 × 10^-10 | 1.62 × 10^-6 |
| 22829 | NLGN4Y | neuroligin 4 Y-linked | -3.967±0.672 | 1.08 × 10^-8 | 1.86 × 10^-5 |
| 27063 | ANKRD1 | ankyrin repeat domain 1 | -2.349±0.445 | 1.52 × 10^-7 | 2.41 × 10^-4 |
| 4633 | MYL2 | myosin light chain 2 | -2.291±0.437 | 1.82 × 10^-7 | 2.67 × 10^-4 |
| 347273 | CAVIN4 | caveolae associated protein 4 | -2.168±0.416 | 2.07 × 10^-7 | 2.82 × 10^-4 |
| 378951 | RBMY1J | RNA binding motif protein Y-linked family 1 member J | -3.824±0.754 | 2.70 × 10^-7 | 3.42 × 10^-4 |
| 1641 | DCX | doublecortin | 3.018±0.662 | 2.24 × 10^-6 | 2.66 × 10^-3 |
| 5940 | RBMY1A1 | RNA binding motif protein Y-linked family 1 member A1 | -3.279±0.765 | 3.71 × 10^-6 | 4.15 × 10^-3 |
| 1264 | CNN1 | calponin 1 | -2.009±0.449 | 8.88 × 10^-6 | 9.39 × 10^-3 |
| 1674 | DES | desmin | -2.452±0.575 | 1.73 × 10^-5 | 1.62 × 10^-2 |
| 728403 | TSPY8 | testis specific protein Y-linked 8 | -3.043±0.768 | 1.71 × 10^-5 | 1.62 × 10^-2 |
| 378949 | RBMY1D | RNA binding motif protein Y-linked family 1 member D | -3.011±0.768 | 1.79 × 10^-5 | 1.62 × 10^-2 |
| 392197 | USP17L7 | ubiquitin specific peptidase 17 like family member 7 | -2.554±0.764 | 5.49 × 10^-5 | 4.75 × 10^-2 |
| 57502 | NLGN4X | neuroligin 4 X-linked | 2.761±0.719 | 6.12 × 10^-5 | 4.92 × 10^-2 |
| 100289087 | TSPY10 | testis specific protein Y-linked 10 | -2.649±0.769 | 6.20 × 10^-5 | 4.92 × 10^-2 |

***Suppl. Table 7: KEGG GOterm hits for Top100 differentially expressed genes LVNC vs resLVNC***.

| ID | Term | Ontology Source | % Associate No. Genes | No. Of Genes | Associated Genes Found |
|---|---|---|---|---|---|
| KEGG:04512 | ECM-receptor interaction | KEGG_13.05.2021 | 6.82 | 6 | COL4A2, ITGA2, ITGA7, ITGB5, LAMB2, THBS1 |
| KEGG:05410 | Hypertrophic cardiomyopathy | KEGG_13.05.2021 | 5.56 | 5 | ITGA2, ITGA7, ITGB5, MYBPC3, TPM1 |
| KEGG:05414 | Dilated cardiomyopathy | KEGG_13.05.2021 | 6.25 | 6 | ADCY6, ITGA2, ITGA7, ITGB5, MYBPC3, TPM1 |

***Suppl. Table 8: KEGG GOterm hits for Top100 differentially expressed genes DCM1 vs resDCM1***.

| ID | Term | Ontology Source | % Associate No. Genes | No. Of Genes | Associated Genes Found |
|---|---|---|---|---|---|
| KEGG:00010 | Glycolysis/ Gluconeogenesis | KEGG_13.05.2021 | 4.48 | 3 | ALDOC, LDHB, PFKM |
| KEGG:03010 | Ribosome | KEGG_13.05.2021 | 8.23 | 13 | RPL10, RPL12, RPL13A, RPL18, RPL19, RPL27, RPL3, RPL7A, RPL8, RPLP0, RPS14, RPS8, RPSA |
| KEGG:05171 | Coronavirus | KEGG_13.05.2021 | 6.03 | 14 | RPL10, RPL12, RPL13A, RPL18, RPL19, RPL27, RPL3, RPL7A, RPL8, RPLP0, RPS14, RPS8, RPSA, STAT3 |
| KEGG:00190 | Oxidative Phosphorylation | KEGG_13.05.2021 | 9.77 | 13 | ATP5F1A, ATP5F1B, ATP5MC2, ATP5PD, ATP8, COX4I1, COX6B1, COX6C, ND4, ND4L, ND5, ND6, NDUFS2 |
| KEGG:04714 | Thermogenesis | KEGG_13.05.2021 | 5.60 | 13 | ATP5F1A, ATP5F1B, ATP5MC2, ATP5PD, ATP8, COX4I1, COX6B1, COX6C, ND4, ND4L, ND5, ND6, NDUFS2 |
| KEGG:05010 | Alzheimer | KEGG_13.05.2021 | 4.07 | 15 | ATP5F1A, ATP5F1B, ATP5MC2, ATP5PD, ATP8, COX4I1, COX6B1, COX6C, CTNNB1, ND4, ND4L, ND5, ND6, NDUFS2, PSMC3 |
| KEGG:05012 | Parkinson | KEGG_13.05.2021 | 6.43 | 16 | ATP5F1A, ATP5F1B, ATP5MC2, ATP5PD, ATP8, COX4I1, COX6B1, COX6C, ND4, ND4L, ND5, ND6, NDUFS2, PSMC3, UBB, UBC |

| | | | | | |
|---|---|---|---|---|---|
| KEGG:05014 | Amyotrophic lateral sclerosis | KEGG_13.05.2021 | 4.12 | 15 | ATP5F1A, ATP5F1B, ATP5MC2, ATP5PD, ATP8, COX4I1, COX6B1, COX6C, ND4, ND4L, ND5, ND6, NDUFS2, PSMC3, VCP |
| KEGG:05016 | Huntington | KEGG_13.05.2021 | 4.9 | 15 | ATP5F1A, ATP5F1B, ATP5MC2, ATP5PD, ATP8, COX4I1, COX6B1, COX6C, ND4, ND4L, ND5, ND6, NDUFS2, POLR2A, PSMC3 |
| KEGG:05020 | Prion disease | KEGG_13.05.2021 | 5.13 | 14 | ATP5F1A, ATP5F1B, ATP5MC2, ATP5PD, ATP8, COX4I1, COX6B1, COX6C, ND4, ND4L, ND5, ND6, NDUFS2, PSMC3 |
| KEGG:05415 | Diabetic cardiomyopathy | KEGG_13.05.2021 | 6.4 | 13 | ATP5F1A, ATP5F1B, ATP5MC2, ATP5PD, ATP8, COX4I1, COX6B1, COX6C, ND4, ND4L, ND5, ND6, NDUFS2 |

**Supplemental figures**

**Suppl. Fig. 1: All iPSC lines show full pluripotency.**

Generated iPSC lines from LVNC-family: LVNC II.3 and from the DCM family: DCM2 III.10, h.RBM1 II.3 and h.RBM2 III.3. h = healthy

**A:** Brightfield and immunofluorescence stainings of iPSC. All iPSC lines are positive for stem cell morphology, ALP activity, and stem cell marker expression OCT4, NANOG, LIN28, and TRA1-60. Brightness and contrast have been enhanced. BF: Brightfield; ALP: alkaline phosphatase.

**B:** All iPSC lines differentiate into cells from all three germ layers stained with the appropriate antibodies against endoderm (AFP: alpha-1-fetoprotein), mesoderm (α-SMA: alpha smooth muscle) and ectoderm (βIII-TUB: tubulin beta 3 class III). Brightness and contrast have been enhanced.

**Suppl. Fig. 2: Genomic integrity and differentiation into iPSC-CM of LVNC- and DCM-affected family members with RBM20 mutations.**

**A:** All iPSC lines retain genetic integrity and a normal karyotype (XX or XY, 46). Exemplary karyotype shown for resLVNC iPSC (XX, 46). Graphs show B Allele Frequency and Log R Ratio for each chromosome. Chr. = chromosome

**B:** Respective Sanger sequencing results of the RBM20 locus exon 9. The zoom visualizes position c.1900-1902, which encodes the amino acid number 634 within the RBM20 gene. Wt-RBM20 encodes C-G-G for p.R634 as an arginine (Arg), whereas LVNC harbors a heterozygous missense mutation C-G/T-G leading to p.R634L (leucine (Leu)) and DCM with C/T-G-G leading to p.R634W (tryptophane (Trp)).

**C:** FLOW cytometry analysis of 60-90 days-old patient and control iPSC-CM stained with cardiac Troponin T (cTNT) antibody. Each dot represents one cardiac differentiation. Undifferentiated iPSC served as negative control.

**D:** QPCR analysis of ventricular marker *MYL2* and atrial marker *NR2F2*. Both genes were normalized to the housekeeper *18s* and subsequently normalized to NR2F2 as a value of one to show the fold change of *MYL2* in relation to the atrial *NR2F2* in every sample. Each dot represents one cardiac differentiation.

**Suppl. Fig. 3: RBM20-dependent mis-localization and *TTN* splicing.**

**A:** RBM20 accumulates in the cytoplasm. Representative immunofluorescence stainings of control-, LVNC-, resLVNC-, DCM1- and resDCM1-CM. Yellow arrows marks exemplary RBM20 cytoplasmatic localization. Antibody staining against RBM20 (yellow) with counterstaining of the nucleus (cyan). Scale bars: 50 µm.

**B:** QPCR analysis of RBM20 splice target *TTN* as *N2B* isoform normalized to *18s*. Data is shown as box plots, whereas every dot represents one differentiation experiment. P-values by Mann-Whitney test.

**C:** The figure shows Principal Component Analysis (PCA) plots generated before differential expression analysis that highlights clusters as expected (patient vs rescue). The PCA plot for LVNC vs DCM1 shows some overlap between the two groups. This suggests that the gene expression profiles of LVNC and DCM1 are more similar to each other than compared to their isogenic controls.

**D, E:** Differential gene expression analysis:

**D:** Volcano plots showing DE genes with key markers labeled in comparison of LVNC vs resLVNC, DCM1 vs resDCM1, and LVNC vs DCM. The -log10 adjusted p-value is plotted against the log2 fold change for each comparison.

**E:** Volcano plots highlighting DE genes that are targets of RBM20 in the same comparison as shown in D. Specific RBM20 target genes are labeled.

DE = Differential (gene) expression, vs = versus

**Suppl. Fig. 4: KEGG GO-term hits (pV<0.05) for the Top100 differentially expressed genes.**

**A:** ResLVNC vs LVNC.

**B:** ResDCM1 vs DCM1.

IA: interaction, HCM: hypertrophic cardiomyopathy, DCM: dilated cardiomyopathy, OxPhos: oxidative phosphorylation, ALS: amyotrophic lateral sclerosis

**Suppl. Fig. 5: Splice graphs with exonic bin usage of LVNC and DCM-specific targets.**

Graphs represent the differential exon expression profiles. The y-axis represents normalized read counts of exons (exon usage), and the x-axis shows individual exons within a gene. The lower panels show the gene structure, with the bars below the x -axis representing exons, and the lines between the bars representing introns. The numbers at the bottom are genomic locations of the gene. Purple bars mark exons with significantly altered usage (FDR-adjusted p-value < 0.05) in the DEX analysis of isogenic versus patient cell lines.

**Suppl. Fig. 6: LVNC- and DCM-CM show differential $Ca^{2+}$ handling pathologies.**

**A:** $Ca^{2+}$ transient decay (TC50) time at basal condition with Fluo-4-AM. LVNC-CM and DCM-CM showed decreased basal decay (TC50) time levels compared to control and rescue lines [number of differentiations/analyzed cells] for control [6/94], LVNC [6/101], resLVNC [3/55], DCM1+2 [7/126] and resDCM1 [3/51]. Data is presented as mean+/- SEM, p-value by Kruskal-Wallis multiple comparisons test with Dunn´s correction. Only significant values are noted.

**B:** No differences in $Ca^{2+}$ SR-load and SR fractional release. Measurements with ratiometric dye Fura-2-AM. P-values were calculated by One-way ANOVA with Dunnett´s correction against control (n.s. = non significant). §P-value was calculated with Mann-Whitney test: patient vs respective isogenic line.

**C:** cAMP changes after Iso treatment among the iPSC-CM lines. Quantification of maximal FRET response to IBMX after Isoprenaline (Iso) stimulation. P-value was calculated by Mann-Whitney test patient vs rescue line but no significances were detected. N.s. = non significant

**Suppl. Fig. 7: Phosphorylation change after Iso treatment and medication of patients.**

**A+B:** Increase in phosphorylation after Iso treatment shown as the fold change from basal to Iso level and normalized to respective isogenic line. P-values by Student´s t-test.

**A:** LVNC-CM exhibit weaker increase in PLN-T17p level after Iso.

**B:** DCM-CM do not differ from resDCM1-CM in Iso response.

**C:** Medication of the patients at the time of biopsies for iPSC generation.

**Suppl. Fig. 8: Characterization of resLVNC-DCM.W.**

**A:** The resLVNC-DCM.W iPSC line is the LVNC patient background with the DCM mutation introduced.

**B:** Brightfield and immunofluorescence stainings of iPSC. The resLVNC-DCM.W-line is positive for stem cell morphology, ALP activity and stem cell markers OCT4, NANOG, LIN28 and TRA1-60. Brightness and contrast has been enhanced. BF: Brightfield; ALP: alkaline phosphatase.

**C:** Molecular karyotyping of resLVNC-DCM.W iPSC showing B Allele Frequency and Log R Ratio for each chromosome. Cells retain a normal karyotype (46, XX). Chr. = chromosome

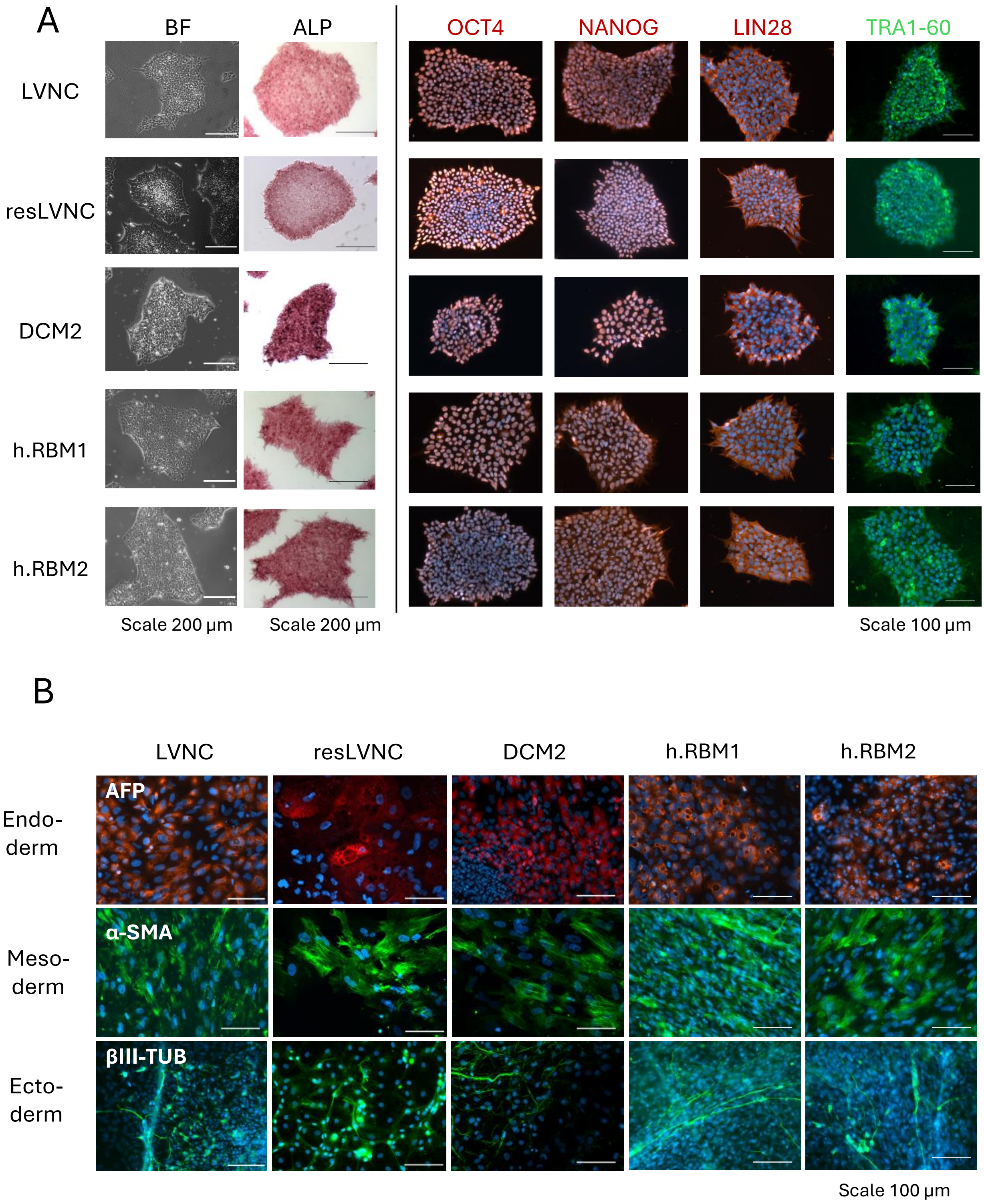


**Suppl. Fig. 1:** All iPSC lines show full pluripotency.

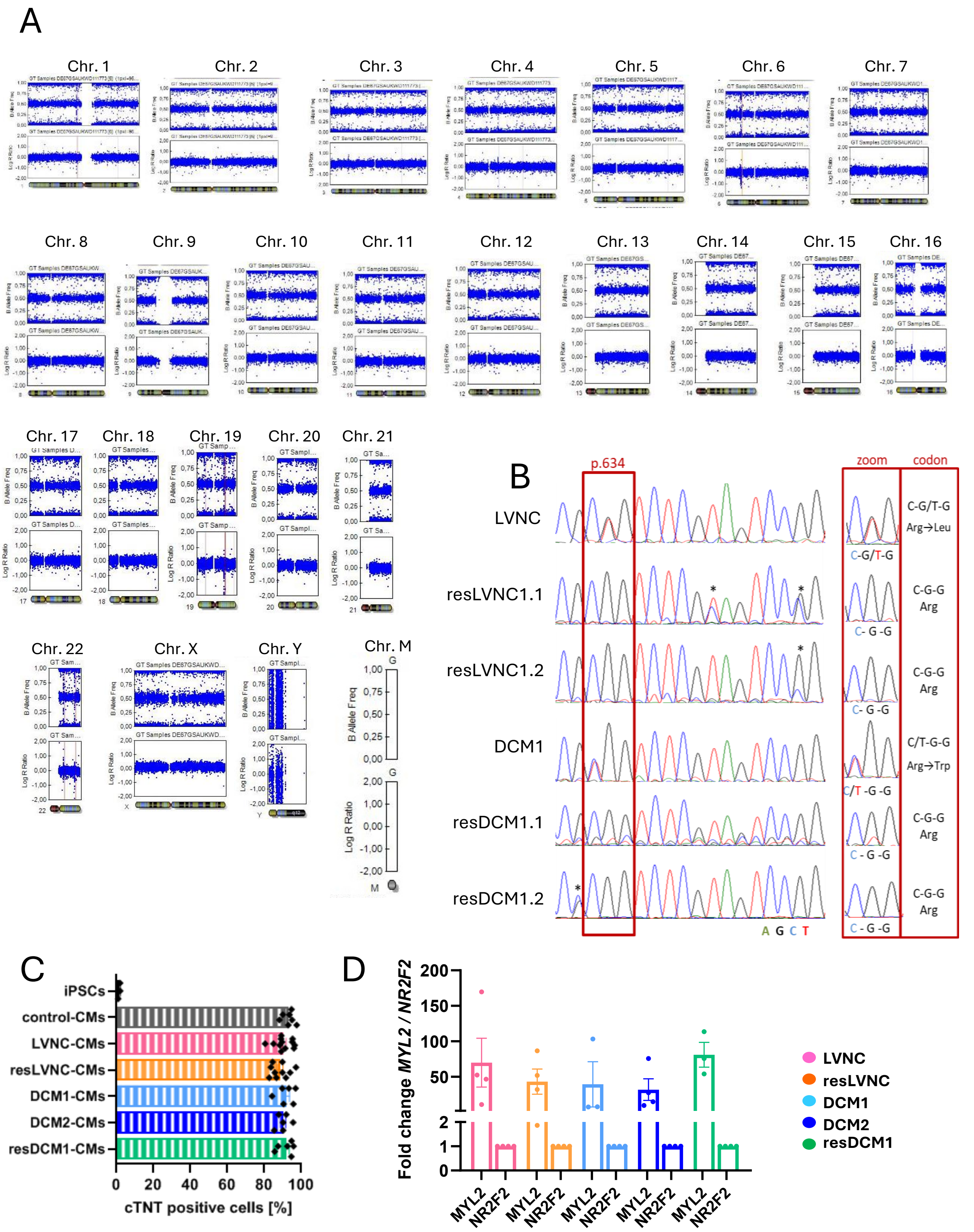


**Suppl. Fig. 2:** Genomic integrity and differentiation into iPSC-CM of LVNC- and DCM-affected family members with RBM20 mutations.

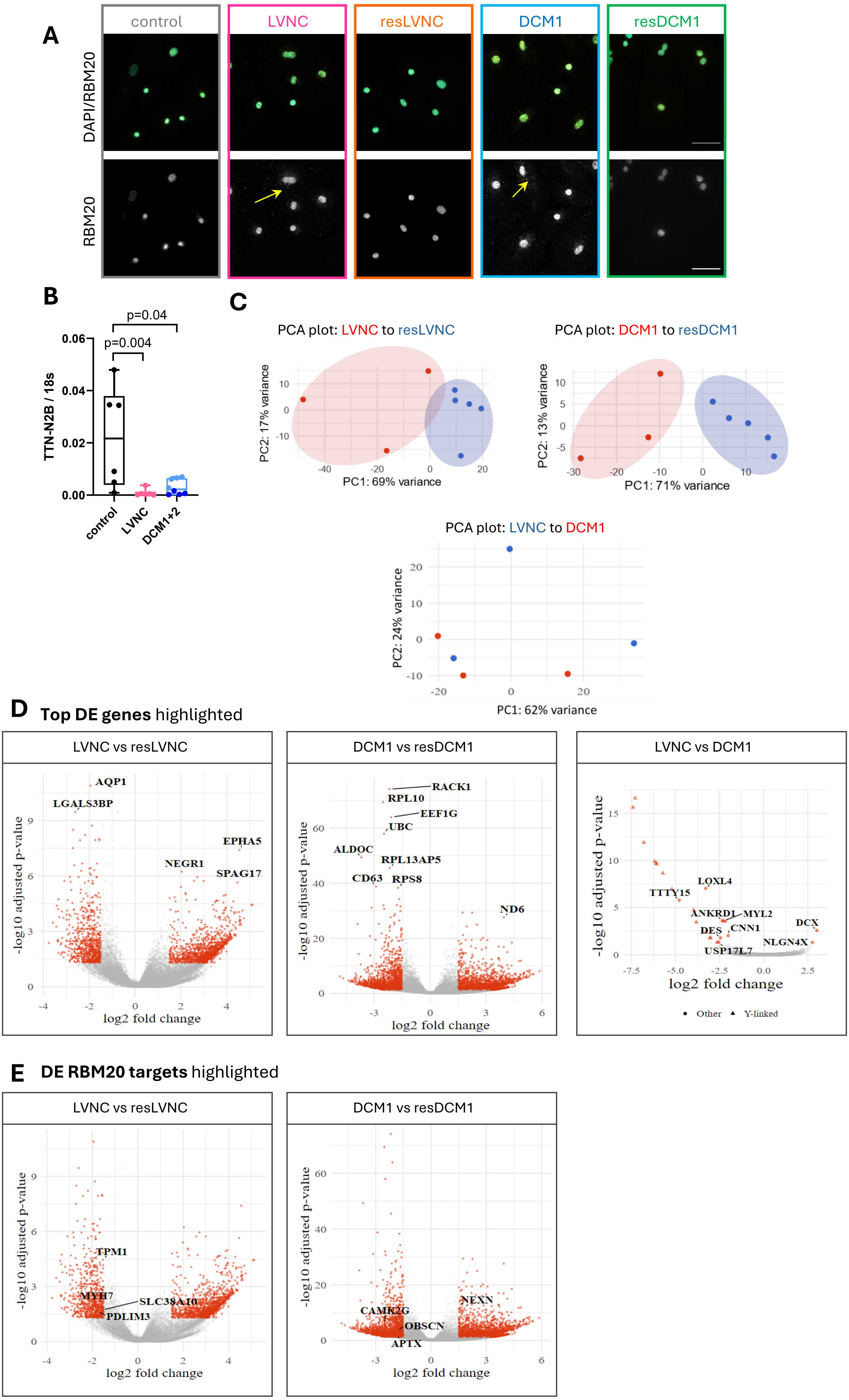


**Suppl. Fig. 3:** RBM20-dependent mis-localization and *TTN* splicing.

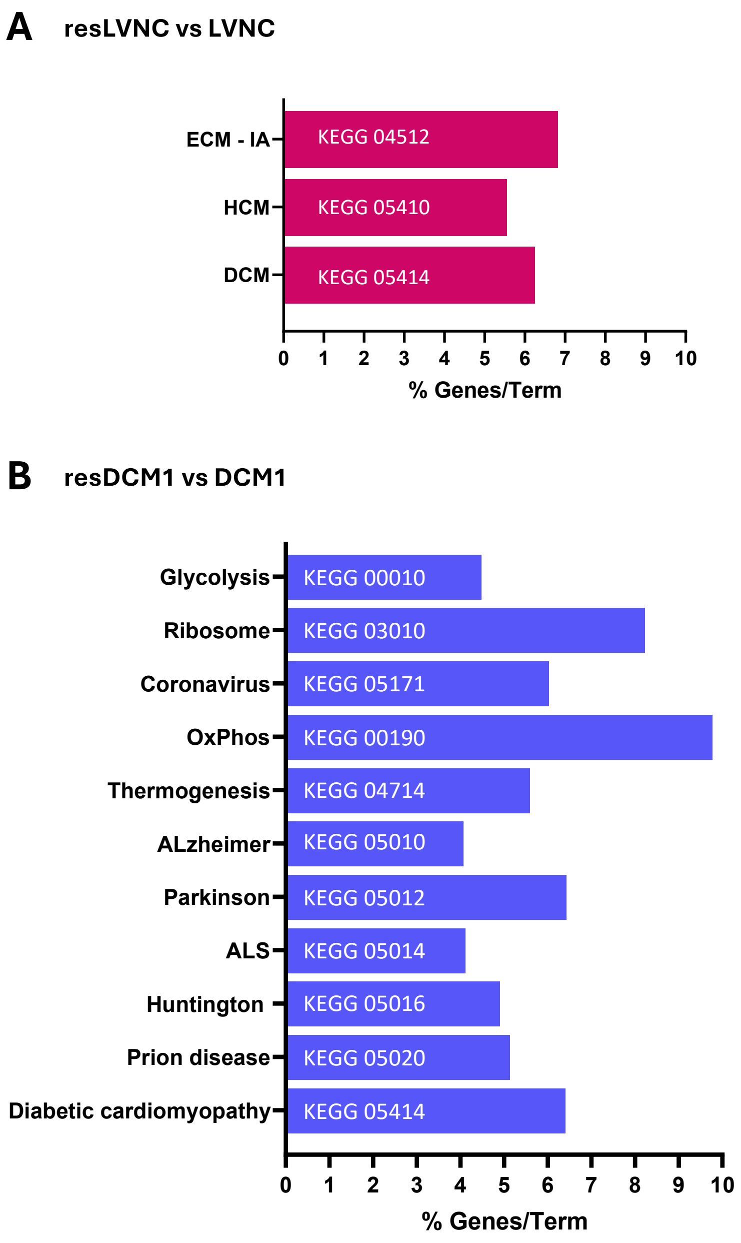


**Suppl. Fig. 4:** KEGG GO-term hits (pV<0.05) for the Top100 differentially expressed genes.

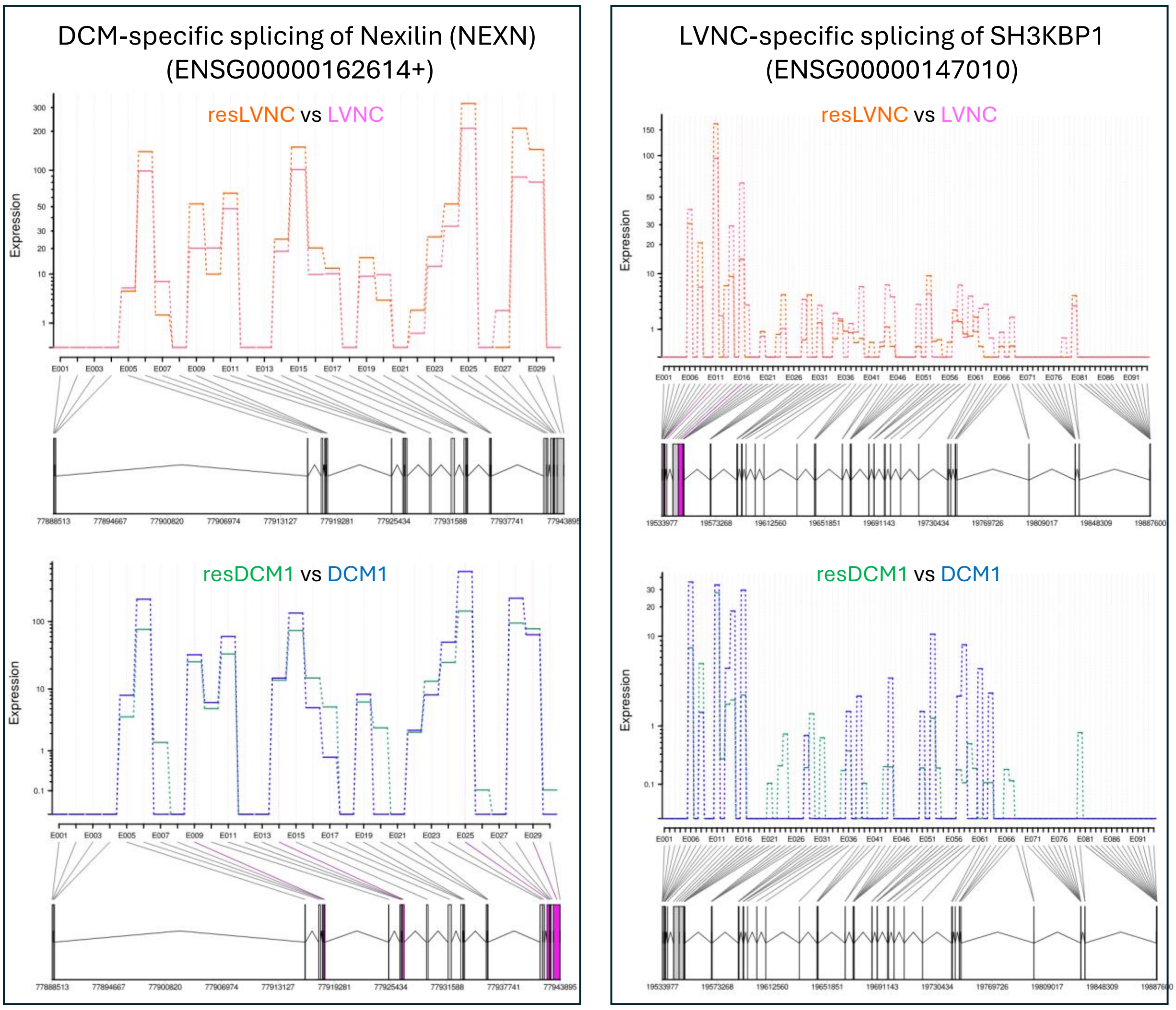


**Suppl. Fig. 5:** Splice graphs with exonic bin usage of LVNC and DCM-specific targets.

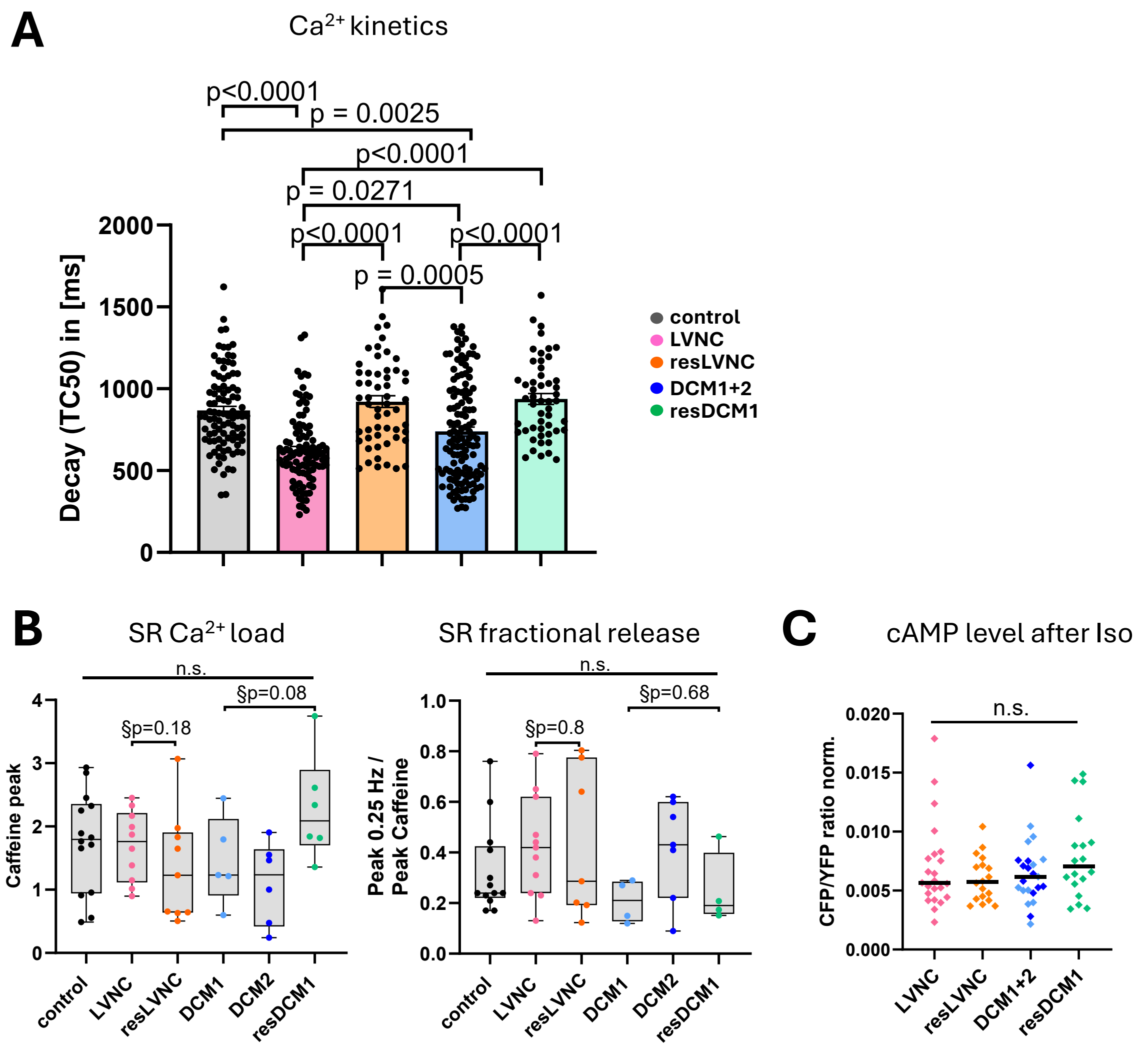


**Suppl. Fig. 6:** LVNC- and DCM-CM show differential $Ca^{2+}$ handling pathologies.

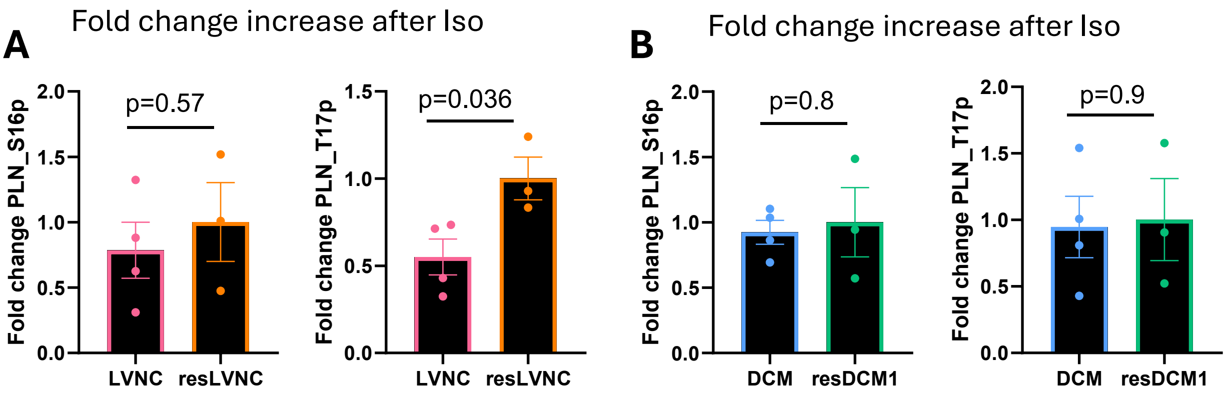


C

| LVNC | DCM1 |
|---|---|
| Ivabradin | Metoprolol |
| Bisprolol | Candesartan |
| Candesartan | Eplerenon |
| Eplerenon | |
| Torasemid | |

**Suppl. Fig. 7:** Phosphorylation change after Iso treatment and medication of patients.

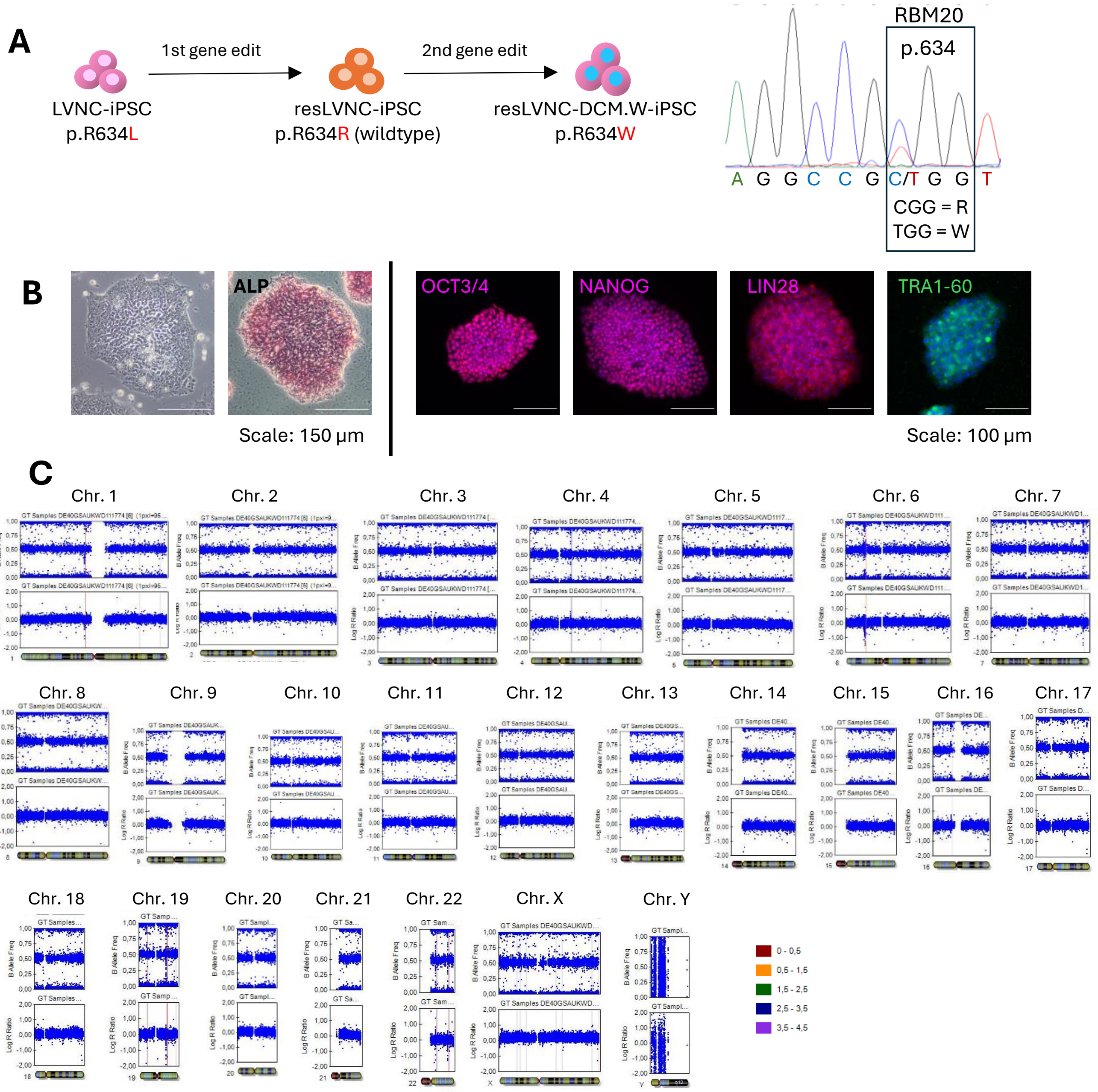


**Suppl. Fig. 8:** Characterization of resLVNC-DCM.W.
